# The evolutionary history and functional specialization of microRNA genes in *Arabidopsis halleri* and *A. lyrata*

**DOI:** 10.1101/2024.05.03.592357

**Authors:** Flavia Pavan, Jacinthe Azevedo Favory, Eléanore Lacoste, Chloé Beaumont, Firas Louis, Christelle Blassiau, Corinne Cruaud, Karine Labadie, Sophie Gallina, Mathieu Genete, Vinod Kumar, Ute Kramer, Rita A. Batista, Claire Patiou, Laurence Debacker, Chloé Ponitzki, Esther Houzé, Eléonore Durand, Jean-Marc Aury, Vincent Castric, Sylvain Legrand

**Affiliations:** Univ. Lille, CNRS, UMR 8198 - Evo-Eco-Paleo, F-59000 Lille, France; Laboratoire Génome et Développement des Plantes, UMR5096 CNRS/UPVD, Perpignan, France; Génomique Métabolique, Genoscope, Institut François Jacob, CEA, CNRS, Univ Evry, Université Paris-Saclay, 91057 Evry, France; Genoscope, Institut de biologie François-Jacob, Commissariat à l’Energie Atomique (CEA), Université Paris-Saclay, Evry, France; Faculty of Biology and Biotechnology, Ruhr University Bochum, D-44801 Bochum, Germany

**Keywords:** Arabidopsis, evolution, microRNA, polymorphism, species-specific genes

## Abstract

MicroRNAs (miRNAs) are a class of small non-coding RNAs that play important regulatory roles in plant genomes. While some miRNA genes are deeply conserved, the majority appear to be species-specific, raising the question of how they emerge and integrate into cellular regulatory networks. To better understand this, we first performed a detailed annotation of miRNA genes in the closely related plants *Arabidopsis halleri* and *A. lyrata* and evaluated their phylogenetic conservation across 87 plant species. We then characterized the process by which newly emerged miRNA genes progressively acquire the properties of “canonical” miRNA genes, in terms of size and stability of the hairpin precursor, loading of their cleavage products into Argonaute proteins, and potential to regulate downstream target genes. Nucleotide polymorphism was lower in the mature miRNA sequence than in the other parts of the hairpin (stem, terminal loop), and the regions of coding sequences targeted by miRNAs also had reduced diversity as compared to their neighboring regions along the genes. These patterns were less pronounced for recently emerged than for evolutionarily conserved miRNA genes, suggesting a weaker selective constraint on the most recent miRNA genes. Our results illustrate the rapid birth-and-death of miRNA genes in plant genomes, and provide a detailed picture of the evolutionary processes by which a small fraction of them eventually integrate into “core” biological processes.

## 1. Introduction

The origins of evolutionary novelties have been a topic of considerable interest in biology (Wagner, 2011). Following Francois Jacob’s (1977) seminal concept of molecular tinkering (Jacob, 1977), the emergence of novel biological functions “from scratch” has long been considered an unlikely evolutionary event. Instead, evolution was believed to proceed through the modification of existing structures (such as protein-coding genes) following various forms of rearrangements such as small or large-scale duplications, fusions or fissions. Recently, however, it has become apparent that new genes do actually arise relatively readily as a result of a variety of processes including pervasive transcription throughout the genome, and the field has moved from “whether” new genes can arise to “how” they arise (Van Oss and Carvunis, 2019). A particularly debated question is whether the newly emerged “proto-genes” become gradually optimized by natural selection and eventually acquire the “canonical” gene-like characteristics (as per the “continuum model”, Carvunis et al., 2012), or whether they result from the immediate apparition of rare “hopeful monsters”, *i.e.* DNA sequences that are already pre-adapted and immediately exhibit gene-like characteristics with essentially no further optimization (as per the “preadaptation model”, Wilson et al., 2017; McLysaght and Guerzoni, 2015). This process has been mostly studied in the particular case of protein-coding genes, and requires the broad-scale comparison of genes of different evolutionary ages that were formed at different times in the past along a phylogeny. However, not all genes are coding for proteins, and the study of the other sorts of genetic elements populating the genome is necessary for a comprehensive understanding of phenotypic evolution.

Regulatory RNAs are an important class of regulators of gene expression, and among them microRNAs (miRNAs) are key post-transcriptional regulators of gene expression in plants, animals, fungi and some viruses (Dexheimer and Cochella, 2020; Nanbo et al., 2021). miRNAs are expressed from genes that do not encode for proteins but are transcribed by RNA polymerase II into primary miRNAs (pri-miRNAs). These pri-miRNAs adopt a hairpin-like structure recognized by DICER-LIKE (DCL) proteins in plants. DCL proteins cleave the pri-miRNA once to generate the pre-miRNA, and a second time to further release the miRNA/miRNA* duplex. The mature miRNA, typically a single 21 nucleotides-long RNA, is loaded into ARGONAUTE1 (AGO1) proteins, forming the RNA-induced silencing complex (RISC) in association with other proteins. The RISC recognizes messenger RNA (mRNA) targets through near-complete sequence complementarity with the mature miRNA, leading to negative regulation through mRNA degradation or translation inhibition (Reviewed in Zhan and Meyers, 2023; Ding and Zhang, 2023).

The recent availability of genome assemblies together with massive small RNA sequencing data has enabled broad-scale comparisons of the repertoire of miRNA genes across a growing number of plant and animal species, albeit with a strong bias toward model organisms. These comparisons revealed striking quantitative variation, with the total number of annotated miRNA genes ranging from just a few dozens to hundreds of miRNA genes per genome (miRBase v22, Kozomara et al., 2019). However, properly interpreting these variations has remained challenging because annotation of miRNA genes in plant and animal genomes is notoriously difficult due to their small size and the abundance of short inverted repeats, the heterogeneity of annotation methods, of the quality of genome assemblies, of molecular methodologies employed for small RNA sequencing, and of tissue types being compared. In spite of these caveats, the data available clearly indicate that, while some miRNAs are deeply conserved, lineage-specific miRNAs are also common, even between closely related species, suggesting a rapid evolutionary dynamics (Fahlgren et al., 2007; Cuperus et al., 2011; Nozawa et al., 2012; Chávez Montes et al., 2014).

The current model posits that “proto-miRNA” genes originate from a variety of sources, including the inverted duplication of protein-coding genes, transposable element-related sequences, or regions of the genome that happened to contain inverted repeats and acquired the ability to be transcribed (reviewed in Cui et al., 2017). However, the abundance of proto-miRNAs relative to canonical miRNAs and the processes by which proto-miRNAs transition into canonical miRNAs have rarely been characterized in detail. Previous studies suggested that the process initially starts from stem-loops exhibiting near perfect complementarity that are the preferred substrate for DCL2, DCL3 or DCL4 proteins, imprecisely generating multiple duplexes of 24-nt-long small interfering RNAs (siRNAs) that are then loaded into AGO4 proteins. As the hairpin structure accumulates mutations over evolutionary time its complementarity is progressively disrupted, facilitating recognition by DCL1 and leading to the precise production of a single 21-nt-long mature miRNA preferentially loaded in AGO1, hence acquiring features of “canonical” miRNA genes (Allen et al., 2004; Voinnet et al., 2009; Baldrich et al., 2018; Pegler et al., 2023). Recent miRNA genes were also suggested to have weaker and more limited spatio-temporal expression territories than the more conserved miRNA genes, and that they also tend to be processed less precisely by DCL proteins leading to the production of a more diverse population of mature miRNAs (Fahlgren et al., 2007, 2010; Ma et al., 2010; Chávez Montes et al., 2014). Young miRNA genes tend to target genes associated with adaptation to local environments (Wen et al., 2016; Bradley et al., 2017), while highly conserved miRNA genes more often target genes involved in crucial cellular processes in plant development and stress responses (Dong et al., 2022). Two models have been proposed for the evolution of miRNA-target interactions. In the “decay model” (Chen and Rajewsky, 2007; Roux et al., 2012), new miRNAs initially have many targets, most of which are deleterious, while only a few are beneficial. Over time, deleterious interactions are removed by natural selection and advantageous interactions are retained. In the “growth model” in contrast, the number of miRNA targets instead increases over the course of evolution (Nozawa et al., 2016). In this model, new miRNAs initially possess few targets, most of which are neutral with only a few being beneficial. This allows the level of expression of the miRNA gene to eventually increase and gradually acquire new targets over the course of evolution. While the target acquisition model has received some support in humans and mice (Chen and Rajewsky, 2007; Roux et al., 2012), the “growth model” has been favored in Drosophila (Nozawa et al., 2016). Hence, the relevance of these models, and the overall evolutionary significance of the newly acquired miRNA genes and their potential regulatory across genomes, has not been widely tested. Finally, young miRNA genes also seem to diverge more rapidly between related species, suggesting weaker functional constraints than that applying to older miRNA genes (Fahlgren et al., 2010; Ma et al., 2010). In *Arabidopsis thaliana,* the binding site within the genes targeted by miRNAs exhibits low polymorphism, indicating strong purifying selection (Ehrenreich and Purugganan, 2008; Smith et al., 2015). However, little is known about the microevolution of the binding site of the genes targeted by recently evolved miRNA genes.

In the genus Arabidopsis, a total of 221 miRNA genes have been annotated in the plant model *A. thaliana* (PmiREN 2.0, Guo et al., 2022b), and the companion papers by Ma et al., (2010) and Fahlgren et al., (2010) identified 154 and 164 miRNA genes, respectively in *A. lyrata*, from which *A. thaliana* diverged about 5 Myrs ago (Koch et al., 2000; Ossowski et al., 2010). These comparisons revealed a series of miRNA genes specific to either species, but given the rapid evolutionary dynamics of miRNA genes such broad-scale phylogenetic comparisons are inherently limited, and the comparison of even more closely related species is needed, as they represent a powerful way to reveal the most recently formed miRNA genes. *A. halleri* diverged from *A. lyrata* only one million years ago (Roux et al., 2011) and is a promising model, but the genome assemblies published for this species are highly fragmented (Briskine et al., 2017; Legrand et al., 2019), and the repertoire of annotated miRNAs is very incomplete (only 18 miRNAs have been deposited in the PmiREN 2.0 database, Guo et al., 2022b).

In this study, we explored the recent evolutionary dynamics of miRNA genes by focusing on *A. halleri* and *A. lyrata*. We first obtained a high-quality chromosome-level reference genome assembly for *A. halleri* and used sRNA-seq data from a variety of accessions to provide the first comprehensive annotation of miRNA genes for this species and followed the same approach to compare them to those in the closely related *A. lyrata* genome. Immunoprecipitation of AGO1 and AGO4 proteins confirmed the validity of the majority of our miRNA gene predictions, including a substantial fraction of those specific to either *A. halleri* or *A. lyrata*, and analysis of the conservation patterns across the Viridiplantae provided a detailed picture of their evolutionary progression along the proto-miRNA - canonical miRNA continuum. Finally, we analyzed whole-genome resequencing data from natural *A. halleri* and *A. lyrata* accessions and showed that the functional constraint varied along the miRNA sequence in a manner that differed according to the evolutionary age of miRNA genes.

## 2. Results

### Reference-level assembly of a *A. halleri* genome

We first produced a chromosome-level reference genome assembly for an individual from Northern France (Auby-1, from the Auby population, 50.40624°N, 3.08265°E) based on a combination of long Oxford Nanopore Technology reads, short Illumina reads and Hi-C data. Briefly, high molecular weight DNA from leaf tissue was extracted and a total of 29 Gb of sequence were obtained using a PromethION (Oxford Nanopore Technology). The 3.32 million reads had a N50 of 18.9 kbp (Supplemental Table S1). The high quality long reads were assembled using NECAT (Chen et al., 2021) and then polished first using all long reads with Racon (Vaser et al., 2017) and Medaka (https://github.com/nanoporetech/medaka) and then using Illumina short-reads with Hapo-G (Aury and Istace, 2021) (Supplemental Table S2). The resulting assembly was composed of 175 contigs and had a cumulative size of 227 Mbp with an N50 of 25.9Mb (Supplemental Table S2). The eight largest contigs covered 90% of the total length and had a size compatible with complete chromosomes (ranging from 22.2 to 31.7 Mbp). The remaining unanchored scaffolds represented only 8.4% of the assembly (Supplemental Figure S1). Hi-C (omni-C) sequencing data were generated to facilitate the chromosome-level assembly and were used to further orientate and anchor contigs to scaffolds (Supplemental Figure S1). We assessed the completeness of the reference genome using BUSCO and found 99.1% complete universal single-copy orthologs, 0.2% fragmented universal single-copy orthologs and 0.7% missing universal single-copy orthologs from the Brassica dataset odb10 (Supplemental Table S2). Overall, the resulting assembly has a sharply higher contiguity than the one published by Legrand et al., (2019) with 18-times less scaffolds and a 93-times higher N50 (Supplemental Table S3).

We used two approaches to annotate protein-coding genes in the genome. First, we aligned the protein sequences of *A. lyrata* and *A. thaliana* against the genome assembly using GeneWise (Birney et al., 2004) to search for homologs. Second, RNA-sequencing data were mapped to the reference genome using Hisat2 (Kim et al., 2019) and assembled by Stringtie (Shumate et al., 2022). Finally, we used Gmove (Dubarry et al., 2016) to combine these two sets of predictions. Overall, a total of 34,721 protein-coding genes were predicted. We used OrthoFinder (Emms and Kelly, 2019) to analyze orthology relationships between the predicted genes of *A. halleri*, *A. lyrata* and *A. thaliana*. After removing orthogroups containing paralogs, we identify 20,306 orthologous genes between *A. halleri* and *A. lyrata*, 13,082 orthologous genes between *A. lyrata* and *A. thaliana* and 13,977 orthologous genes between *A. halleri* and *A. thaliana*.

### Annotation of the miRNA genes in the *A. halleri* Auby1 individual

To obtain a comprehensive set of miRNA genes in the *A. halleri* reference genome, we first generated ultra-deep small RNA sequencing (sRNA-seq) data from two tissues (leaves and a mix of flower buds at different stages of development) of the accession used to produce the reference genome (Auby-1). We obtained a total of 206 and 159 million Illumina reads for the two sRNA-seq libraries (leaves and buds, respectively) (Supplemental Table S4). To enhance our ability to annotate miRNA genes, we combined predictions from miRkwood (Guigon et al., 2019) and Shortstack (Johnson et al., 2016), two algorithms that are adapted for plant genomes. While Shorstack is more conservative and predicts fewer miRNAs, miRkwood includes less reliable miRNA predictions but still with a majority of the miRNAs predicted in *A. thaliana* loaded in AGO1 or AGO4, which is considered high-level evidence for their regulatory potential (Guigon et al., 2019). Overall, after merging the predictions from the two tissues, we obtained a total of 332 predicted miRNA genes in the *A. halleri* reference genome (Supplemental Table S4).

To investigate whether sequencing depth could be a limiting factor for the discovery of miRNA genes, we randomly sub-sampled sequencing reads from the library with the highest number of reads (the one obtained from leaves, comprising 206 million reads), and newly predicted the miRNA genes using the exact same procedure in ten independent replicates for each sample size. We observed that a depth of 165 million reads is required to predict 90% of the total set of miRNA genes (Figure 1a), and observed no clear plateau of the number of predicted miRNA genes, indicating that even such a high sequencing depth remains a limiting factor, and that more miRNA genes with low abundance remain to be discovered. However, we note a clear inflection of the saturation curve once the first 86 miRNA genes have been discovered, suggesting that a limited set of miRNAs with relatively high abundance can already be revealed with a lower sequencing depth (around 20 million reads, as is classically done in many sRNA sequencing experiments).

**Figure 1.**
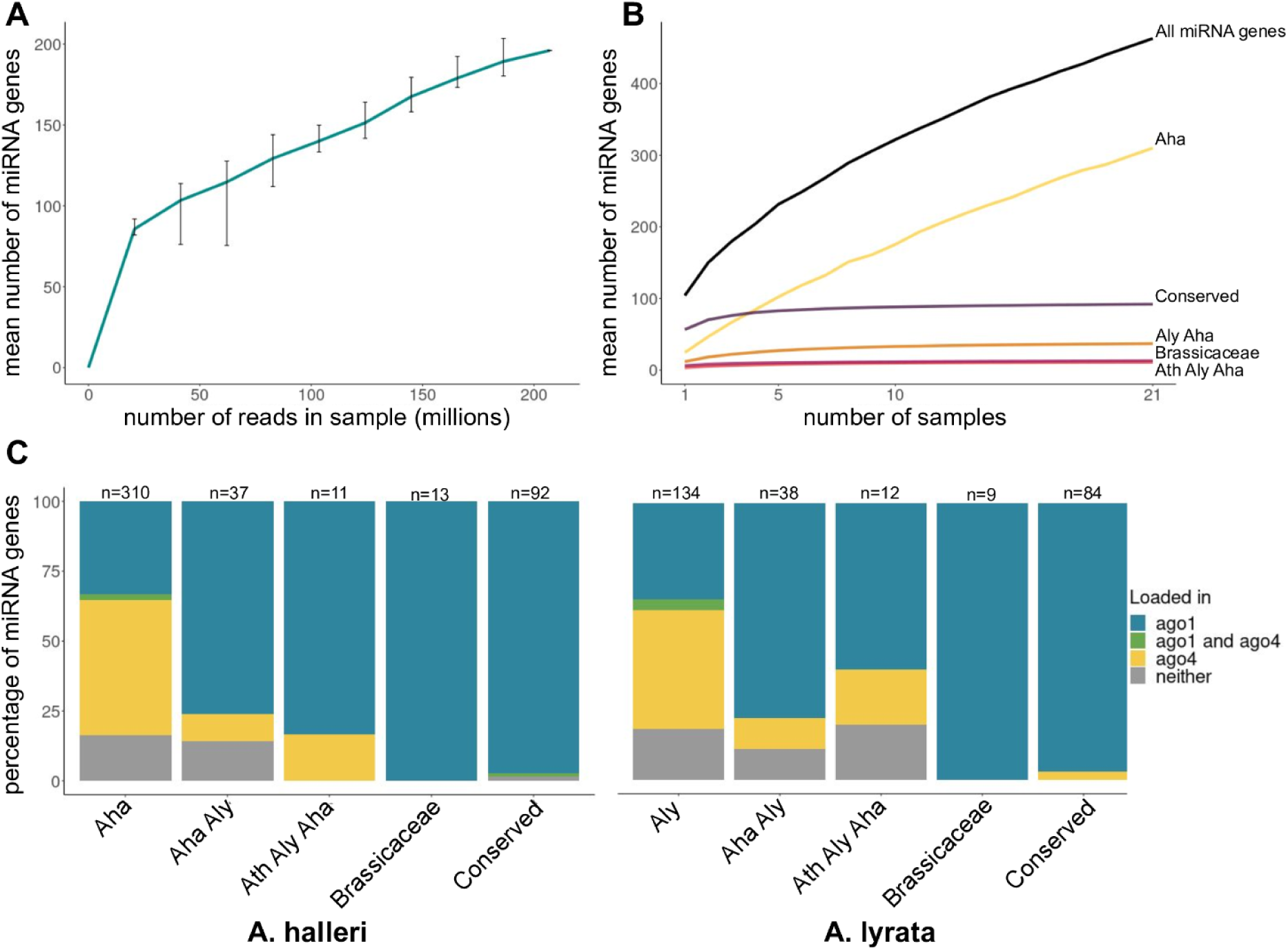
Annotation of miRNA gene repertoires. (a) Sequencing depth saturation curve for the most deeply sequenced accession (*A. halleri* Auby-1 accession, leaves). Bars represent the 95% confidence interval. (b) Sample saturation curve for miRNA gene repertoires in *A. halleri* according to their conservation. (c) Percentage of miRNA genes loaded in AGO1 and/or AGO4 proteins out of 314 and 147 miRNA genes expressed in the input fractions in *A. halleri* and *A. lyrata*.

### Core and accessory miRNA genes in the *A. halleri* and *A. lyrata* reference genomes

To evaluate the variation of the repertoire of miRNA genes, we then aligned sRNA-seq data that we either generated ourselves (*n* = 6 libraries) or retrieved from the Sequence Read Archive (SRA) at the NCBI (*n* = 13 libraries) onto the *A. halleri* reference genome. For *A. lyrata*, we used the recently updated reference genome (accession MN47, Kolesnikova et al., 2023) and aligned reads from *n* = 3 sRNA-seq libraries that we generated and *n* = 10 sRNA-seq publicly available libraries. These data originate from a diversity of geographical accessions, plant tissues (leaves, buds and roots), developmental stages, sample preparation (such as True-seq, Nextflex Small RNA-Seq, SOLiD Total RNA-Seq, ION total RNA-seq), sequencing methods (SOLID, PROTON, Illumina) and sequencing depths (from two to 206 million reads) (Supplemental Table S4). Given this heterogeneity, the results are expected to buffer the inherent technical biases associated with individual sRNA sequencing experiments (Wright et al., 2019). Our analysis predicted between 46 and 267 miRNA genes per sample (Supplemental Table S4). After merging the predictions across samples, we identified a total of 463 and 276 miRNA genes in *A. halleri* and *A. lyrata*, respectively (Supplemental Data Set S1). The higher number detected in *A. halleri* is expected because of the larger number of sequencing datasets analyzed. Because a given miRNA precursor could produce different mature miRNA molecules in different accessions, these miRNA genes together resulted in a total of 678 and 521 mature miRNAs in *A. halleri* and *A. lyrata*, respectively (*i.e.* on average, a miRNA gene produced 1.5 and 1.9 mature miRNAs across all accessions in *A. halleri* and *A. lyrata*) (Supplemental Data Set S1; Supplemental Data Set S2). About a third of these miRNA genes were predicted by both softwares (287 in *A. halleri* and 87 in *A. lyrata*), while 145 and 176 genes were unique to miRkwood and 31 and 13 were unique to Shortstack in *A. halleri* and *A. lyrata,* respectively. The higher number of predictions made by miRkwood is in line with Li et al., (2021), who showed that miRkwood is able to predict substantially more miRNAs than other plant miRNA prediction tools.

### Completeness of the repertoires

While some miRNA genes were broadly shared and predicted in at least 80% of the samples (8.6% and 9.8% in *A. halleri* and *A. lyrata*), a large proportion was private to a single sample (52.3% and 42.8% in *A. halleri* and *A. lyrata*) (Figure 2a, b). Hence, the number of “core” miRNA genes was relatively limited as compared to the accessory miRNome, noting that different individuals from the same species can carry or express different miRNA genes because of genetic or environmental variation. To evaluate the completeness of the set of miRNA genes we predicted in the *A. halleri* and *A. lyrata* genomes, we performed a saturation analysis by randomly subsampling within the 21 and 13 individual samples from each species. We performed 1,000 replicates for each sample size and evaluated how the number of miRNA predictions increased with the number of samples upon which they are based. For *A. halleri,* we found that 18 of the 21 samples were needed to reach 90% of the total number of predictions (Figure 1b). Similarly, in *A. lyrata* 8 of the 11 samples needed to be included to reach 90% of the total number of predictions (Supplemental Figure S2). Hence, it is clear that the repertoire of miRNA genes in these two species was not saturated and was limited by the number of accessions that have been sequenced so far. In particular, our results show that analyses based on a single sequencing experiment in a single reference accession (as commonly performed) are likely underestimating the number of miRNA genes in a species by at least an order of magnitude. Altogether, our results suggest that our ability to discover miRNA genes remains limited both by the number of accessions and the sequencing depth.

**Figure 2:**
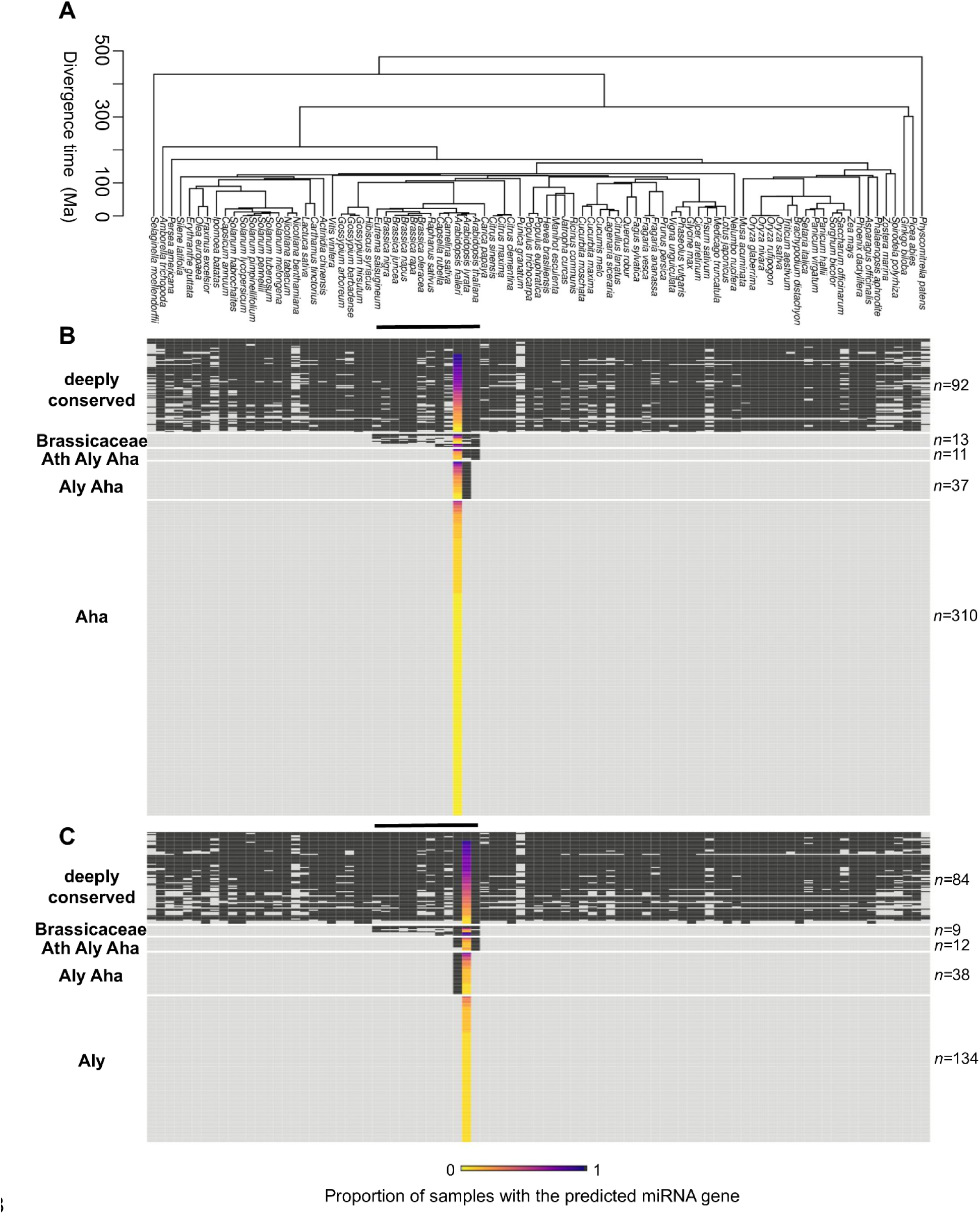
The majority of miRNA genes is either deeply conserved or species-specific. (a) Phylogenetic tree based on TimeTree v.5 of 87 Viridiplantae species present in the PmiREN database. Phylogenetic families are shown in supplementary table S6. The black bars indicate the Brassicaceae family. Overview of the miRNA gene conservation (b) in *A. halleri* and (c) in *A. lyrata*. Each line corresponds to one miRNA gene, and species are represented in columns. Black squares indicate the presence of an homolog/ortholog, and gray squares its absence in the corresponding species. The number of miRNA genes in the five groups of conservation are indicated on the left part of the figure (1: deeply conserved, 2: shared with the Brassicaceae family, 3: shared with the Arabidopsis family, 4: shared between *A. halleri* and *A. lyrata,* 5: species-specific). The proportion of accessions in which the miRNA gene was predicted is indicated by the colored bars from yellow (unique sample) to black (all samples).

### A majority of miRNA predictions are validated by AGO-IP

We then performed immunoprecipitation of AGO1 and AGO4 proteins to provide formal experimental validation of our predictions. To broaden the set of miRNA genes we could discover, we analyzed three tissues (leaves, buds and roots) from a pool of six *A. halleri* or *A. lyrata* individuals per populations (Auby, France and I9, Italy for *A. halleri* and Plech, Germany for *A. lyrata*), and sequenced the small RNAs associated with these proteins as well as the input material (total cellular fraction). Out of the total set of miRNA genes predicted above, 314 and 147 were present in the *A. halleri* and *A. lyrata* input samples. A large majority of these predicted miRNA genes (88.2% and 83.4%, respectively) produced mature miRNAs associated with either AGO1 or AGO4 proteins (Figure 1c). Consistent with previous findings (Mi et al., 2008), the sRNAs loaded in AGO1 were predominantly 21-nucleotides-long with a 5’ uridine (38.6% in *A. halleri* and 33.05% in *A. lyrata*), while the sRNAs loaded in AGO4 were predominantly 24-nucleotides-long with a 5’ adenosine (56.2% in *A. halleri* and 60.6% in *A. lyrata*) (Supplemental Figure S3). Therefore, our bioinformatic annotation strategy identifies *bona fide* miRNAs with a substantial number of canonical miRNA genes, including a large number of those that are accession-specific.

### A minority of miRNA genes are conserved at a large phylogenetic scale

To evaluate the evolutionary age of miRNA genes, we combined three different strategies at increasingly divergent phylogenetic scales. First, we aimed at a fine-scale comparison between *A. halleri* and *A. lyrata.* To do this, we considered miRNA genes as syntenic if their hairpin sequences were reciprocal best-hits and if they were flanked by syntenic genes. Second, we took advantage of the availability of assembled genomes and sRNA sequencing experiments in eleven Brassicaceae species to apply the exact same discovery pipeline we used in *A. halleri* and *A. lyrata.* Details of the genome assemblies and sRNA-seq experiments included are reported in Supplemental Table S5. Note that because of divergent genome structures and variable quality of the genome assemblies we did not attempt to recover synteny relationships for this phylogenetic level. Finally, we extended the analysis to the broad set of Viridiplantae species included in the PmiREN 1.0 (Supplemental Table S6). database, which was constructed by uniformly processing sRNA sequencing datasets and uses a recent set of criteria to identify miRNA genes (Guo et al., 2020). Because precursor sequences can diverge rapidly, we aligned the mature miRNAs predicted in *A. halleri* and *A. lyrata* (rather than full precursor sequences) to the mature miRNAs predicted in the 87 species of the database (Figure 2a), and considered mature miRNAs as homologous if they shared ⩾ 85% sequence similarity. Combining the results of the three analyses, we observed very similar patterns of conservation in both species (Figure 2b, c). We defined five groups of conservation for which we associated an age based on the divergence time in the phylogeny: 1) deeply conserved miRNAs shared by very distant species. This category represents 20% (*n* = 92) and 30% (*n* = 84) of the predicted miRNAs genes in *A. halleri* and *A. lyrata*, respectively (Figure 2b, c), and includes well-studied miRNA families such as miR156/miR157, miR166/miR161, miR169 and miR395. 2) miRNAs shared across the Brassicaceae family. This category represents 3% (*n* = 13) in *A. halleri* and 3% (*n* = 9) in *A. lyrata* (Figure 2b, c), and also includes some well-studied miRNA families such as miR158, miR845, miR400. 3) miRNAs shared between *A. thaliana*, *A. halleri* and *A. lyrata*. This category represents 2% (*n* = 11) in *A. halleri* and 4% (*n* = 12) in *A. lyrata* (Figure 2b, c), and also includes some well-studied miRNA families such as miR822, miR823 and miR842. 4) miRNAs shared only between *A. halleri* and *A. lyrata*. This category represents 8% (*n* = 37) in *A. halleri* and 14% (*n* = 38) in *A. lyrata*, including previously annotated families such as miR3433 and miR3443. 5) the *A. halleri*- specific miRNAs represent 67% of the *A. halleri* repertoire (*n* = 310) and the *A. lyrata*-specific miRNAs represents 49% of the *A. lyrata* repertoire (*n* = 134) (Figure 2b, c). Based on the divergence time between these two closely related species (Roux et al., 2011), we estimate that this last category of miRNAs appeared at most one million years ago. Overall, in both species we found that the vast majority of annotated miRNAs were either broadly conserved or species-specific, with only a small fraction of miRNAs showing intermediate levels of phylogenetic conservation.

### Natural variation of the repertoire of deeply conserved and species-specific miRNAs

We then determined how the set of miRNA genes in each group of conservation varied with the number of samples included in the analysis. The species-specific miRNA genes tended to be detected in a smaller number of samples (8.2% and 10.8% of the samples in *A. halleri* and *A. lyrata,* respectively) than the deeply conserved genes (detected in 61.8% and 59.6% of the sample in *A. halleri* and *A. lyrata,* Figure 2b,c). Specifically, in *A. halleri*, 90% of the total number of predictions of the most deeply conserved miRNA genes were already annotated with only five of the 21 samples. Similarly, only six samples were needed to annotate 90% of the total number of miRNA genes shared within the Brassicaceae family, respectively. In contrast, up to nine and 14 samples were needed to annotate miRNA genes shared with *A. lyrata* and the *A. halleri*-specific genes (Figure 1b). Similarly, in *A. lyrata*, only five and three individuals were needed to identify 90% of the deeply conserved and the miRNA genes shared with the Brassicaceae family, while up to eight and nine individuals were needed for the miRNA genes shared with *A. halleri* and the *A. lyrata*-specific miRNA genes (Supplemental Figure S2). Overall, these results indicate that our analysis of multiple samples probably represents a comprehensive set of the deeply conserved miRNA genes, while the repertoire of species-specific miRNA genes is not saturated even with a large number of samples. Hence, including more samples would probably mostly increase the number of species-specific miRNA genes.

### How young miRNAs become canonical miRNAs

Based on our evaluation of the evolutionary age of miRNA genes, we then sought to characterize how the more recent miRNAs genes differ from the more ancient ones. For this analysis, we merged the orthologous miRNA genes between *A. halleri* and *A. lyrata,* for which we took the average of each character value, *i.e* reads abundance, length, stability, processing precision. Our dataset was composed of 97 deeply conserved miRNA genes, 14 genes shared within the Brassicaceae family, 14 genes shared between *A. thaliana*, *A. halleri* and *A. lyrata,* 38 genes shared between *A. halleri* and *A. lyrata*, and 444 *A. halleri*- or *A. lyrata*-specific miRNA genes. We used linear regression models to evaluate how a series of molecular properties evolved with age of the miRNA genes. The mean normalized level of expression of the miRNA genes increased from 74.4 reads per kilobase million mapped reads (RPKM) for the species-specific miRNA genes to 278.9 RPKM for the most deeply conserved (adjusted R²=0.28; p-value=2.66e-55) (Figure 3a; Supplemental Figure S5). Similarly, expression of the mature miRNA increased from 2.8 to 35.9 RPM (adjusted R²=0.34; p-value=1.58e-70) (Supplemental Figure S4). These results are consistent with previous studies that showed that conserved miRNA genes tend to be expressed broadly at higher levels than the more recent miRNA genes (Cuperus et al., 2011). We note that in spite of this general trend, there is a strong variance within categories (as evidenced by the low adjusted R ^2^), and some of the most recent miRNA genes could still be expressed quite substantially, at levels comparable to those of some of the most conserved miRNAs.

**Figure 3:**
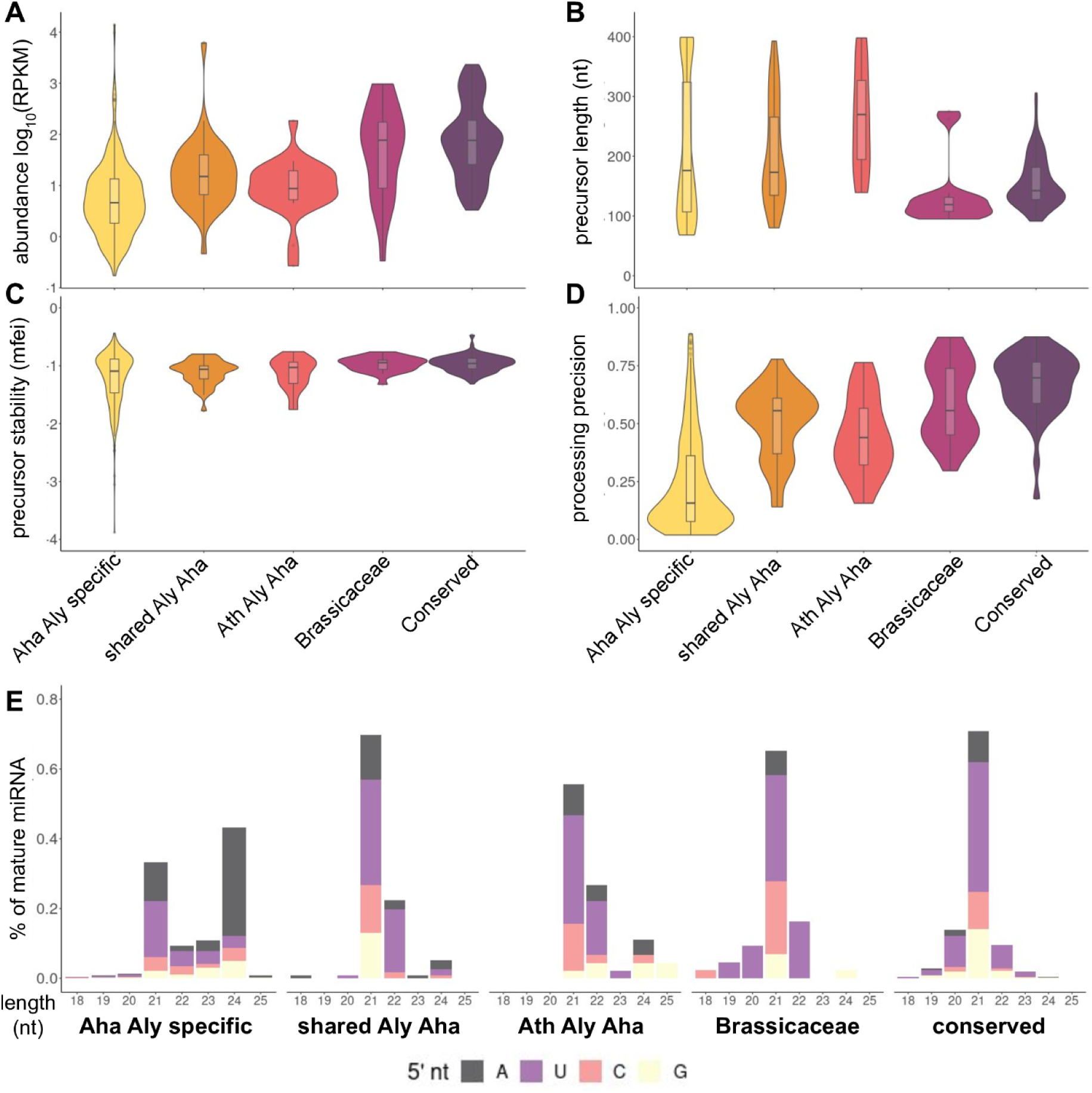
The characteristics of miRNA genes evolve slightly in the course of evolution. The orthologous genes between *A. halleri* and *A. lyrata* have been merged and categorized in five groups of conservation: the deeply conserved miRNA genes, those shared with the Brassicaceae family, those shared among *A. thaliana*, *A. halleri* and *A. lyrata*, those shared exclusively between *A. halleri* and *A. lyrata*, and the species-specific miRNA genes. (a) Expression level of the miRNA genes. (b) Length of the hairpin produced by the miRNA gene. (c) Hairpin stability, estimated using Minimum Free Energy Index (MFEI) calculation (d) DCL processing precision. (e) Mature miRNA size distribution and nature of the first 5’ nucleotide.

Second, we analyzed the evolution of the size, stability and processing precision of the hairpins produced by the miRNA genes. We found that the average hairpin length tended to decrease over the course of evolution, with relatively long hairpins for species-specific and Arabidopsis-specific miRNA genes (mean size of 213 nt and 261 nt, respectively), but a shorter mean size of only 155 nt for the deeply conserved miRNAs (adjusted R²= 0.05; p-value=1.57e-10) (Figure 3b; Supplemental Figure S5). We then estimated the minimal free energy index (MFEI) of each hairpin as an indicator of stability, for which a score far from zero indicates high stability. The average MFEI increased from the species-specific (−1.21) to the deeply conserved miRNA genes (−0.96; adjusted R²=0.07; p-value=3.86e-13) (Figure 3c; Supplemental Figure S5), corresponding to a decrease of the stability of the hairpin structure as miRNAs became more ancient. The hairpin structure, in particular the presence of bulges, is an important factor for cleavage by DCL proteins (Bologna et al., 2013). Following Ma et al., (2010), we defined the DCL processing precision of each miRNA gene as the abundance of mature miRNA sequences divided by the abundance of all the reads mapping to the hairpin. A score close to one indicates a high processing precision by DCL, while a score close to zero indicates an imprecise processing. The average processing precision increased from the species-specific miRNA genes (0.24) to the deeply conserved miRNA genes (0.67; adjusted R²=0.38; p- value=4.24e-78) (Figure 3d; Supplemental Figure S5). Altogether, our results show that over the course of evolution, the hairpin produced by miRNA gene decreases in length, becomes more unstable and is processed more precisely by DCL proteins. Third, we examined the size and 5’ nucleotide of miRNAs, as these features are known to be important for miRNA biogenesis and functions (Mi et al., 2008). The proportion of 21-nucleotides miRNAs with a uridine as the first 5’ nucleotide increased from the species-specific miRNAs (15%) to the deeply conserved (37%), while the proportion of 24-nucleotides miRNAs with an adenosine as the first 5’ nucleotide decreased from 32% (species-specifics) to 0.3% (deeply conserved) (Figure 3e). AGO1 proteins select mainly 21-nucleotides miRNAs with a 5’ uridine, while AGO4 proteins select mainly 24-nucleotides miRNAs with a 5’ adenosine (Mi et al., 2008), and accordingly we found that the vast majority of the conserved miRNAs were loaded in AGO1. This was especially true for the most conserved miRNAs (71/72, 99% and 66/68, 97% in *A. halleri* and *A. lyrata* respectively), but also for the miRNAs shared across the Brassicaceae family (100% for both species, 8/8 and 7/7 in *A. halleri* and *A. lyrata* respectively), those shared across the Arabidopsis genus (5/6, 83% and 3/5, 60% in *A. halleri* and *A. lyrata*) and those shared by *A. halleri* and *A. lyrata* miRNAs (16/21, 89% and 14/18, 78% in *A. lyrata*). A substantial proportion of the *A. halleri*- and of the *A. lyrata*-specific miRNAs (68/207, 33% and 17/49, 35% respectively) were also loaded in AGO1 (Figure 1c). Loading into AGO4 followed the opposite trend, as 99 of the 207 (48%) *A. halleri*-specific and 21 of the 49 (43%) *A. lyrata*-specific miRNAs were found in the AGO4 fraction. This proportion decreased rapidly as miRNA genes became older, with only 2/21 (9%) and 2/18 (11%) for miRNA shared by *A. halleri* and *A. lyrata,* respectively, 1/6 (17%) and 1/5 (20%) for miRNAs shared across the three Arabidopsis species, in *A. halleri* and in *A. lyrata*, respectively. None of the deeply conserved miRNAs in *A. halleri* and only two of the 68 deeply conserved miRNAs in *A. lyrata* (3%) were associated with AGO4 (Figure 1c). Finally, dual loading in both AGO1 and AGO4 was relatively rare, with only 7/207 of the *A. halleri*-specific, 2/49 of the *A. lyrata*-specific and 2/72 of the deeply conserved miRNAs in *A. halleri* being almost equally loaded in AGO1 and AGO4 (Figure 1c). Hence, our results provide a clear picture, where miRNAs produced by nearly all ancient miRNA genes are almost exclusively loaded in AGO1, while miRNAs produced by the very young miRNA genes are mainly loaded in AGO4 and a substantial proportion in AGO1. Thus, in spite of their limited conservation, a substantial proportion of these species-specific miRNAs may already have some regulatory potential.

### The number of essential targets increases over the course of evolution

To gain insight into the integration of miRNA genes in gene regulatory networks, we predicted the targets in the coding sequences (CDS) across the genome for each miRNA gene using Targetfinder (Bo and Wang, 2005), and first evaluated the evolution of their number according to the age of the miRNA gene. The number of predicted targeted genes per miRNA gene was positively correlated with its age (adjusted R²=0.17; p-value=3.09e-32) (Supplemental Figure S6), increasing from species-specific miRNA genes (0.87 targets on average per miRNA) to the most deeply conserved (5.42 targets on average per miRNA) (Figure 4a). Second, we determined the essentiality of the genes targeted by miRNAs using three proxies as in Legrand et al., (2019): 1) the size of the gene family (single-copy genes are predicted to be more essential due to the lack of functional redundancy); 2) the *k*_A_/*k*_S_ ratio calculated from orthologous genes between *A. halleri, A. lyrata* and *A. thaliana*, for which lower values are expected for more essential genes; 3) the presence of loss-of-function (LOF) phenotype in *A. thaliana* mutant*s* (Lloyd and Meinke, 2012). After merging the orthologous miRNA genes, our dataset was composed of 262 genes targeted by the most deeply conserved miRNA genes, 40 by the miRNA genes shared across the Brassicaceae family, 17 by the miRNA genes shared between *A. thaliana*, *A. halleri* and *A. lyrata,* 129 by the miRNA genes shared between *A. halleri* and *A. lyrata* and 150 by the species-specific miRNA genes. The k_A_/k_S_ ratios calculated from *A. halleri, A. lyrata* and *A. thaliana* divergence were negatively correlated with age of the miRNA gene (adjusted R²=0.02; p-value=0.03), with a mean k_A_/k_S_ of 0.22 and 0.33 for the genes targeted by species-specific miRNAs and miRNAs shared between *A. halleri* and *A. lyrata* respectively, and a lower value (k_A_/k_S_=0.18) for the genes targeted by the deeply conserved miRNAs (Figure 4b; Supplemental Figure S6). The average gene family size of the genes targeted was negatively correlated with age of the miRNA genes (adjusted R²=0.01; p-value=0.003), decreasing from 1.47 and 5.01 for the genes targeted by species-specific and Brassicaceae-specific miRNAs to 1.38 for those targeted by deeply conserved miRNAs (Figure 4c; Supplemental Figure S6). The frequency of target genes with a LOF phenotype was correlated with age of the miRNA gene (p-value=0.009). However, the frequency of LOF genes initially decreased slightly (from 0.087 for the genes targeted by the species-specific genes, close to the genomic average, to 0.066 for the genes targeted by miRNAs shared by *A. halleri* and *A. lyrata*), but then increased sharply to 0.179 for those targeted by deeply conserved miRNAs (Figure 4d). Overall, our results indicate that the number of miRNA-target interactions increases over the course of evolution, along with the proportion of interactions involving essential genes.

**Figure 4:**
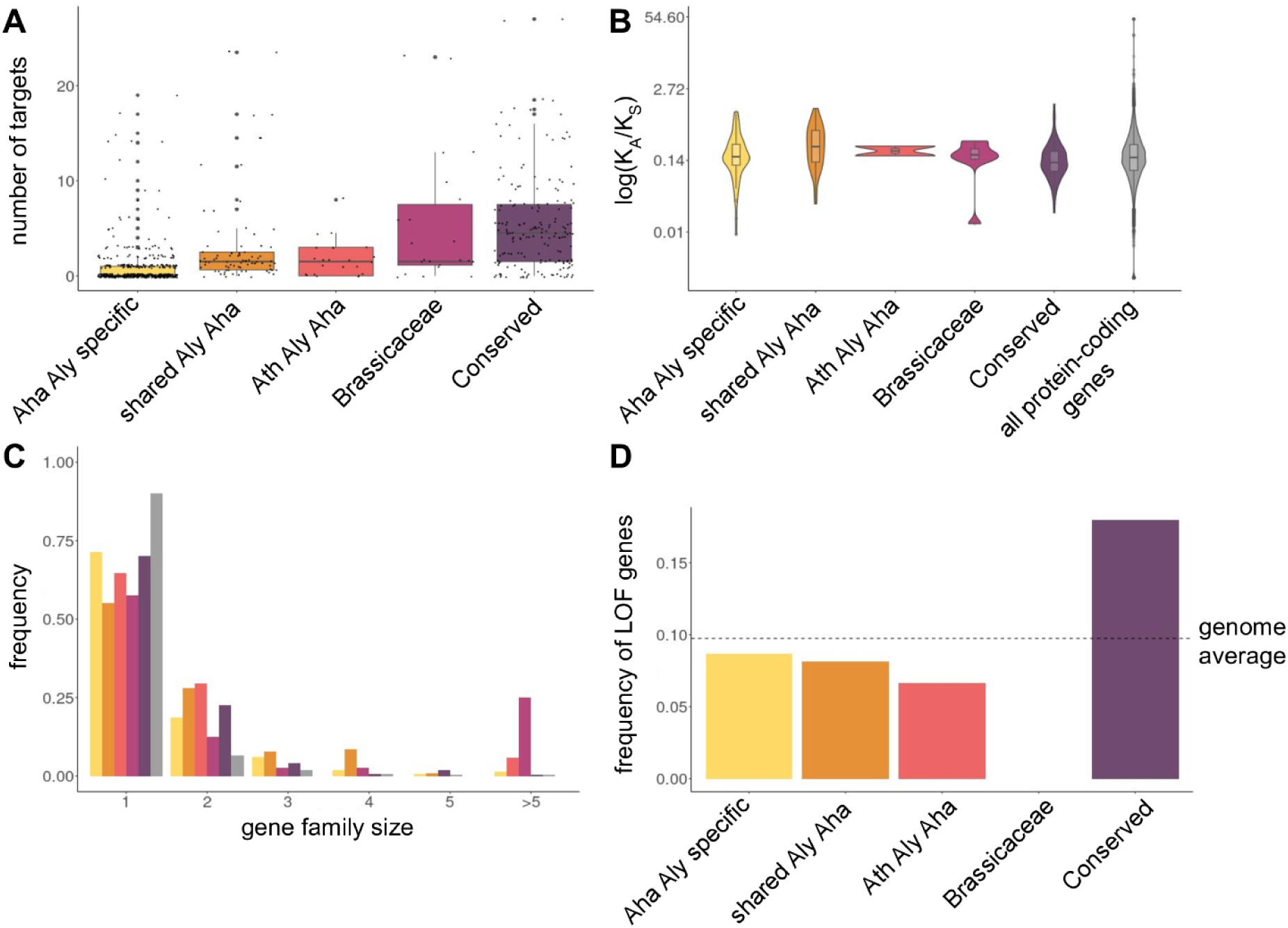
The old miRNA genes have more essential targets than the young ones. (a) Number of targets per miRNA gene according to its age. (b) k_A_/k_S_ ratios of targeted genes calculated from *A. halleri, A. lyrata* and *A. thaliana* divergence. (c) Frequency of the targeted gene family size. (d) Frequency of LOF phenotype genes in the miRNA genes targets. The frequency of LOF phenotype genes in all the genes present in the species is indicated by the dashed line.

### Functional constraint on the miRNA/miRNA* duplex over the course of evolution

To determine whether certain parts of the hairpin were more constrained by natural selection than others, we investigated nucleotide polymorphism of the 276 *A. lyrata* miRNA genes in 100 *A. lyrata* individuals from natural populations that we either newly sequenced or retrieved from published datasets. We determined the level of nucleotide polymorphism (*π*) for each part of the miRNA hairpins, including the miRNA, the miRNA*, the rest of the stem, the loop, as well as 200 bp of upstream and downstream flanking sequences (Figure 5a). Polymorphism of the miRNA/miRNA* duplex showed a decrease of about 53% in *A. lyrata* compared to the rest of the precursor (*π=*0.0062 vs. 0.0134), suggesting high selective constraint (Supplemental Figure S7). Strikingly, polymorphism of the duplex in the deeply conserved miRNA genes (mean *π* of 0.0062) was equivalent to the polymorphism of the 0-fold degenerate positions of protein-coding genes (mean *π* of 0.0047 for both species), suggesting that this part of the precursor evolves under considerable selective constraint (Figure 5b). The overall level of polymorphism of the hairpin decreased from the species-specific (mean *π* of 0.0153) to the deeply conserved miRNA genes (0.0076) (Figure 5b). Polymorphism of the species-specific miRNA genes was similar to the polymorphism of the 4-fold degenerate positions across the genome (mean *π* of 0.0150). Thus, our results suggest that, collectively, the youngest miRNA genes tend to evolve close to neutrality, although we note that this conclusion does not preclude the possibility that some of them may be involved in the control of important biological functions. In contrast, the selective constraint on the more deeply conserved miRNAs is considerable, with levels of polymorphism of the miRNA/miRNA* duplex even lower than those of the most strongly constrained sites of protein-coding sequences.

**Figure 5:**
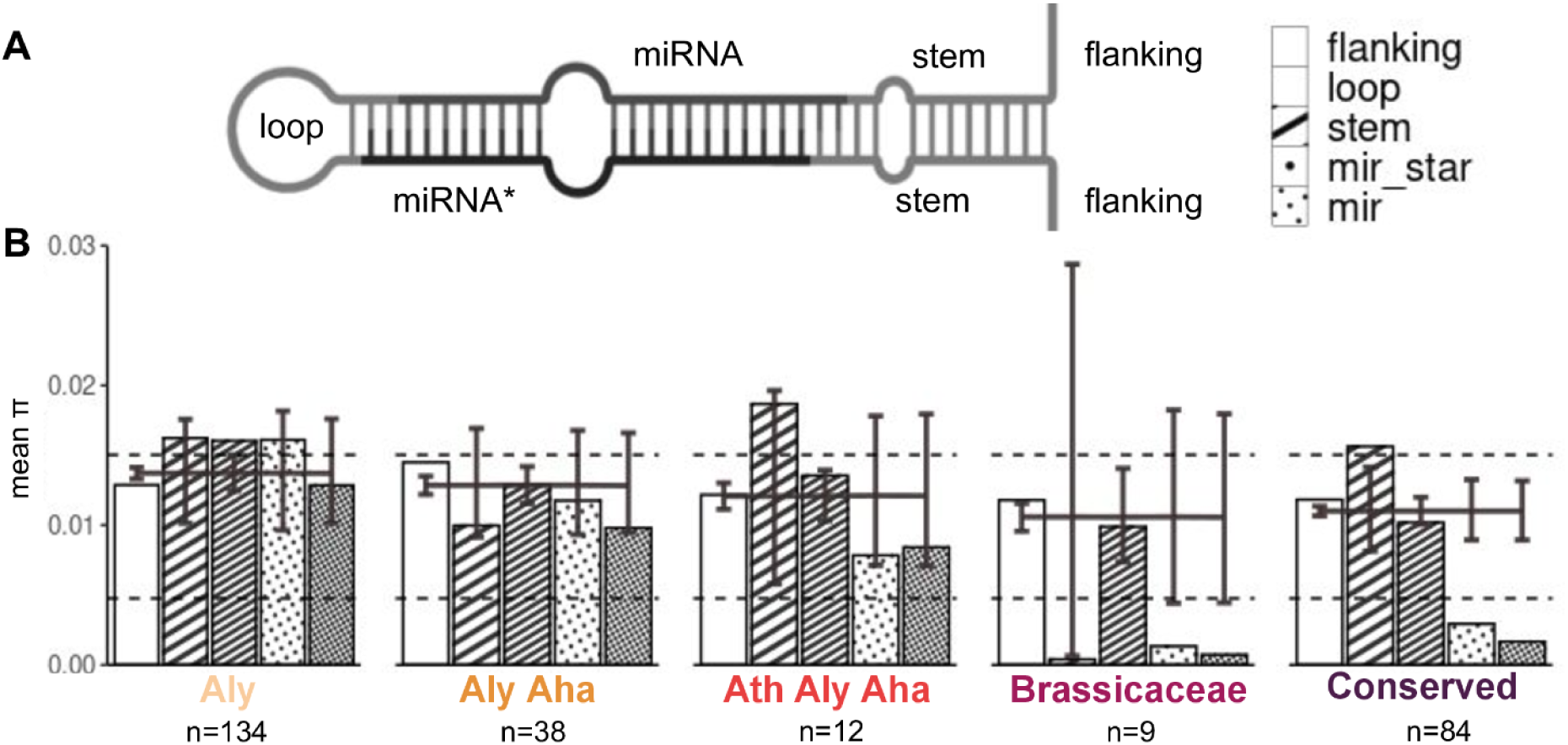
Selective constraints increase over the course of the evolution of miRNA genes. (a) Description of the miRNA hairpin regions with the upstream and downstream flanking regions (200 bp each). (b) *A. lyrata* average nucleotide diversity in the different parts of the miRNA hairpins. The dashed lines represent the mean π values for the 0 fold (lower line) and 4 fold (upper line) degenerate positions of all protein-coding genes of the genome. The bars represent the 95% confidence interval obtained by random permutation of nucleotides for 1,000 simulations.

### Natural selection on the miRNA binding sites

We then asked whether the targeting by miRNAs could represent a detectable functional constraint along the coding sequence of their target genes. To test this hypothesis, we compared the polymorphism of 1,042 predicted binding sites in *A. lyrata* with that of their 300 bp upstream and downstream flanking regions along the target mRNAs. We observed slightly lower polymorphism of the binding site (average *π* 0.0090 in *A. lyrata*) as compared to the flanking regions (average *π* 0.0098 in *A. lyrata*), *i.e.* a 8.8% reduction, suggesting that the presence of the miRNA binding site represents a detectable selective constraint on the CDS in addition to the original constraint of coding for a specific set of amino-acids (Figure 6, Figure S8).

**Figure 6:**
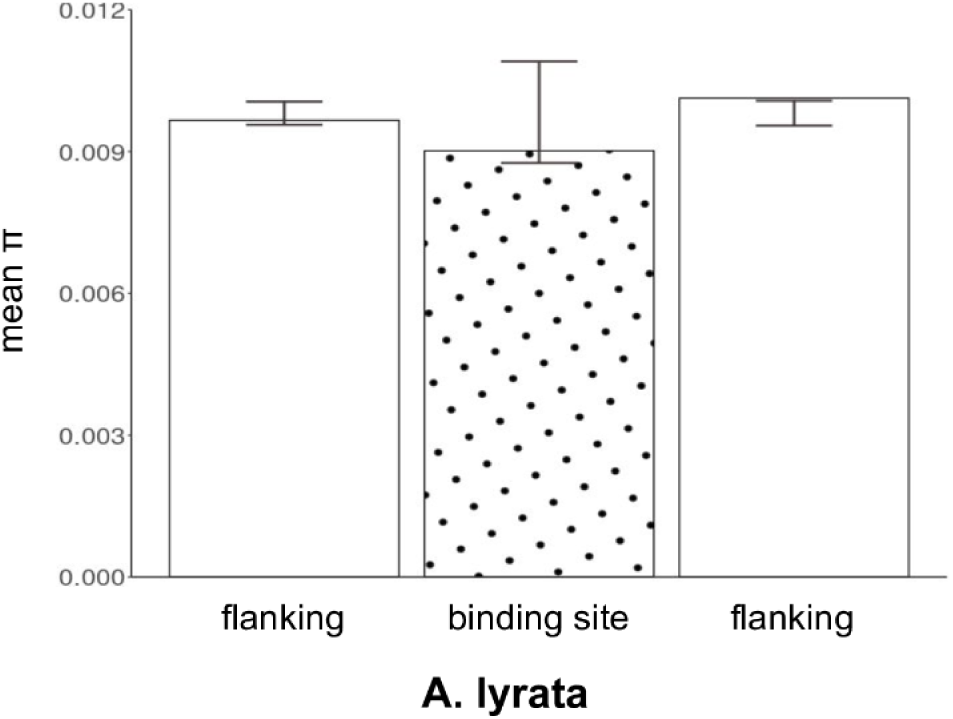
**miRNA binding sites** (dotted bar) **have lower** average nucleotide diversity in *A. lyrata* than their upstream and downstream flanking regions (300 bp each, white bars). The whiskers represent the 95% confidence interval under the assumption of a random distribution of polymorphisms along the concatenated sequence and were obtained by random permutation of nucleotides for 1,000 simulations.

## 3. Discussion

### Challenges in the identification of miRNAs in plant genomes

Proto-miRNAs have been proposed to emerge relatively readily, but studying their emergence and evolution has remained challenging because this requires the comparison of well-assembled and well-annotated closely related genomes, and high-quality deep sRNA sequencing data. Our deep miRNA annotation of the closely related *A. halleri* and *A. lyrata* genomes revealed both the long-term conservation and important evolutionary lability of these genetic elements. Identifying the complete set of miRNA genes in a species remains challenging for at least two reasons. First, in line with Ma et al., (2010) and Cuperus et al., (2011), we find that evolutionarily young miRNA genes tend to be expressed at low levels, and Chávez Montes et al., (2014) suggested that their expression territories might be limited in space and in time. Our results show that in spite of our extensive sequencing of a diversity of samples from diverse environmental conditions, tissues, or accessions of origin, the discovery of new miRNA genes is not yet exhausted in *A. halleri* and *A. lyrata*. Specifically, we observed a relatively limited “core” miRNAome, and the majority of miRNA genes belong to the “accessory” miRNAome, found in a single or a few samples only. To achieve an even more comprehensive annotation, it will now be necessary to take into account the genomic variation among accessions by moving away from the alignment of sRNA-seq reads onto a single reference genome, and assembling individual genomes across a diverse set of natural accessions. While we took advantage of published data from a diversity of sources to maximize the number of accessions and environmental conditions, an important limitation is that we did not control for these factors. For a more detailed analysis it will also be necessary to compare miRNA annotations across accessions cultivated under a common garden environment. The second challenge is that annotation of miRNA genes relies on a set of criteria that have remained debated (reviewed in Axtell and Meyers, 2018). Here, we show that even though young miRNA genes tend to exhibit non-canonical features and can thus be hard to distinguish from false positives, in line with Guiguon et al., (2019) we observed that a vast majority of the predicted miRNA genes were experimentally validated by AGO1- and AGO4-IP, including a substantial fraction of the most evolutionarily recent ones. We conclude that the criteria we used for miRNA annotation were relatively stringent, and that the regulatory potential conferred by loading into AGO1 and/or AGO4 seems to be acquired rapidly, at least for some of them.

Fahlgren et al., (2010) compared miRNAs in *A. lyrata* and *A. thaliana* and estimated that 18% of them were *A. lyrata*-specific and 22% were *A. thaliana* specific. Here, with our deeper annotation, we found an even higher proportion, with up to 67% *A. halleri-* and 49% *A. lyrata*-specific miRNA genes. This difference illustrates the increased sequencing power over the last decade, and the effect of our strategy to multiply the number of accessions. In addition, Fahlgren et al., (2010) estimated that 134 miRNA genes were shared exclusively between *A. lyrata* and *A. thaliana*. Here, by including a much larger set of outgroup species (87 species in the Pmiren database), we restricted this set to only 11 or 12 miRNA genes specific to the Arabidopsis genus (depending on whether they were seen specifically in *A. halleri* or *A. lyrata*). This further illustrates that the vast majority of miRNA genes are either deeply conserved or species-specific, with very few showing intermediate levels of conservation. This small number might still be an overestimation, since the set of mature miRNA sequences for some species in the PmiREN database is probably incomplete (*e.g. Nicotiana benthamiana n*=73, *Punica granatum n*=33 or *Pisum sativum n*=51), possibly explaining the lack of homologs detected in the deeply conserved group for some species.

### The evolutionary history of miRNA genes

A large proportion of the miRNA genes we identified are species-specific and have emerged recently, providing unprecedented power to explore the early steps of their evolution. Collectively, our results largely support the verbal model of emergence of miRNA genes proposed earlier (Allen et al., 2004; Voinnet et al., 2009; Baldrich et al., 2018; Pegler et al., 2023), whereby young miRNA genes start from near-perfect and relatively long hairpins, whose length and stability decrease by an accumulation of mutations creating bulges over the course of evolution together with a decrease of the diversity in size of the mature miRNA population, while the overall expression level and processing precision of the hairpin increases. Our findings parallel the observations made in the context of the *de novo* birth of protein-coding genes (Carnuvis et al., 2012; Wilson et al., 2017). Just like a large number of ORFs can be identified in a genome, we also identified a very large number of potential candidates being tested by natural selection, with possibly neutral or deleterious effects initially. Then only a very small fraction are retained over the long run, and eventually control essential cellular functions. The question of whether the miRNA genes that eventually become fixed have been slowly optimized by natural selection from imperfect progenitors, or rather represent “hopeful monsters” that were immediately beneficial when they arose is difficult to address directly. Yet, we note that the variance of molecular features among the group of the most recent miRNA genes is very large, so their distribution largely overlaps with that of the canonical miRNA genes. Hence, our results are consistent with the idea that at least some of them may already exhibit features allowing them to function as efficiently as the highly conserved canonical miRNA genes.

### Origin of young miRNA genes

Previous studies in rice (Zhang et al. 2011), wheat (Poretti et al., 2019; Crescente et al. 2022) and more generally in angiosperms (Guo et al. 2022a) suggested that transposons are an important source of origin of new miRNA genes. In contrast, Fahlgren et al. (2010) and Nozawa et al. (2012) suggested that a large proportion of miRNA genes in *A. thaliana* emerged from protein-coding genes. These differences observed between plant species could be due to variation of the genomic content of transposable elements in these species, or related to the age of the miRNA genes used in these studies. In our study, we observe a high proportion of young miRNA genes that produce 24 nt sRNAs and are loaded into AGO4, indicating their possible integration into RNA-directed DNA Methylation (RdDM) pathway. One of the main roles of the RdDM pathway is the stable silencing of transposons (Erdmann et al. 2020), thus it is tempting to speculate that the young miRNA genes whose products are loaded in AGO4 could originate from transposons and may participate in their regulation, as was documented e.g. by Borges et al. (2018). An interesting next step will be to formally evaluate the contribution of the different possible progenitor sequences that could have given rise to the vast repertoire of new miRNA genes we uncovered.

### Integration of young miRNA genes in the regulatory network

Although there are examples of young miRNA genes having important functional roles (Wen et al., 2016; Bradley et al., 2017), our results suggest most of them are unlikely to have essential biological functions, and are rapidly lost by genetic drift, mutation or natural selection (Fahlgren et al., 2010; Ma et al., 2010; Nozawa et al., 2012; Smith et al., 2015). Here, we found that the expression level of young miRNA genes was low and their miRNA/miRNA* duplex evolved largely neutrally, suggesting that these genes may not have a significant effect on the cell or the individual. On the other hand, some miRNA genes are deeply conserved and the question of how a new miRNA integrates the functional regulatory network without impairing the fitness of the individual is still debated. Chen and Rajewsky, (2007) argued that in humans young miRNA genes have many targets that appear at random in the genome, few of which are neutral or advantageous and many of which are slightly deleterious and will be lost. In contrast, Nozawa et al., (2016) argued that young miRNA genes have only few targets, most of which are neutral, and only a small fraction of which are beneficial. Neutral miRNA-targets interactions are rapidly lost through drift mutations, while beneficial ones are conserved under purifying selection. During this period, the expression level of the miRNA can increase to enable efficient suppression of its important targets and the miRNA may also acquire new targets because the chances of forming pairs with mRNAs is higher when it is more highly expressed itself (Nozawa et al., 2016). We observed that the young miRNA genes have fewer essential targets than older ones, supporting the “growth model”. Nonetheless, we found that the proportion of these interactions involving essential genes decreased before increasing again in the deeply conserved genes. This trend could result from natural selection initially removing deleterious interactions as the expression level of the miRNA gene increases.

A striking result of our analysis is the reduced nucleotide diversity of the miRNA binding site along the mRNA sequence. However, the extent of the reduction we observed is a lot weaker than that observed in *A. thaliana* by Ehrenreich and Purugganan, (2008). This study focused on miRNA binding sites that were validated by experimental data, so are probably enriched for the interactions with the strongest magnitude of regulatory effect. In addition, the annotation of miRNAs in this study relied on more limited data, and so were also enriched for the “low hanging”, most highly expressed miRNA genes that are easier to detect. It would be interesting to extend our analysis to evaluate the effect of the choice of miRNA-target interactions on the magnitude of the reduced diversity within the target sites.

### Evolutionary significance of new miRNA genes

It is clear from our results that not all miRNA genes in a genome have the same evolutionary age. Some have been present for extended periods of time, while others emerged very recently. While it is clear that the most conserved miRNA genes fulfill essential biological functions, the evolutionary significance of the species-specific miRNA genes is harder to establish. This difficulty parallels that encountered for other genomic elements or cellular features. For instance, the evolution of long non-coding RNAs has been hotly debated. While key roles have been documented for some, (e.g. Statello et al., 2021), overall they seem to have little to no actual evolutionary importance, and most of them are largely dispensable (Goudarzi et al., 2019). Similarly, while alternative splicing is now recognized as a widespread phenomenon, the fraction of alternative splicing events with actual adaptive role is possibly low, and the variation of this feature among species is best explained by the drift-barrier hypothesis (Benitière et al. 2023; Lynch 2007). Here, even though the species-specific miRNA are not conserved, we cannot exclude that some have important biological functions. One example of non-conserved miRNA genes obviously fulfilling an important biological function is given by the sRNA precursors controlling dominance interactions between self-incompatibility alleles in Arabidopsis (Durand et al., 2014). Similarly, sRNAs determining the patterns of adaptation to the local environments encountered by specific accessions would also not be expected to show strong conservation. Given the large number of new miRNA being tested by natural selection at a given time, it is possible that non-conserved miRNA may play an important role in the rapid adaptation of plants to changing environments. At the same time, it is also possible that the majority of species-specific miRNA genes may in fact be neutral, as suggested by their low number of predicted targets and the fact that the proportion of LOF genes among their predicted target genes closely matches that of a random draw across the genome. To achieve a better understanding of the origin of new miRNA genes, it will now be necessary to investigate the molecular nature of their potential progenitors across the genome. In addition, their actual regulatory impact is currently hard to measure, and speculation can only be made on the basis of very indirect evidence. Designing experiments to determine whether at least some of them actually have the capacity to regulate their predicted target genes will be a challenging, yet fascinating next step.

## 4. Materials and methods

### Plant material

*A. halleri* and *A. lyrata* plants were gathered from natural populations (see Supplementary table S6) and subsequently cultivated in standard greenhouse conditions. This cultivation aimed to produce leaves and buds for DNA and RNA extractions. For argonaute immunoprecipitation experiments, cuttings from six *A. halleri* Auby, ten *A. halleri* I9 and six *A. lyrata* Plech individuals were cultivated in hydroponic conditions in a growth medium composed of 1 mM Ca(NO _3_)_2_, 0.5 mM MgSO_4_, 3 mM KNO_3_, 0.5 mM NH_4_H_2_PO_4_, 0.1 μM CuSO_4_, 0.1 μM NaCl, 1 μM KCl, 2 µM MnSO_4_, 25 μM H_3_BO_3_, 0.1 μM (NH_4_)_6_Mo7O_24_, 20 μM FeEDDHA, and 1 μM (*A. lyrata*) or 10 μM (*A. halleri*) ZnSO_4_. The pH of the solution was maintained at 5.0 using MES acid buffer (2 mM). Roots were collected after six weeks.

### *A. halleri* reference genome

#### High-molecular-weight DNA extraction, PromethION library preparation and sequencing

Two grams of fresh leaves were collected and flash-frozen. High molecular weight genomic DNA was extracted as described in (Belser et al., 2018 and Vacherie et al., 2022). For Nanopore library preparation, the smallest genomic DNA fragments were first eliminated using the Short Read Eliminator Kit (Pacific Biosciences, Menlo Park, CA, USA). Starting with 1 µg of genomic DNA, libraries were then prepared according to the protocol « 1D Native barcoding genomic DNA (with EXP-NBD104 and SQK-LSK109) » provided by Oxford Nanopore Technologies (Oxford Nanopore Technologies Ltd, Oxford, UK), with some minor exceptions (increased incubation times for enzymatic steps and purification on beads). All the DNA recovered after the ligation of the barcode step was pooled with the DNA of three other samples before the final adaptor ligation. Each library, containing a total of four barcoded samples, was loaded on a R9.4.1 PromethION flow cell. In order to maintain the translocation speed, flow cells were refueled with 250µl Flush Buffer when necessary. Reads were basecalled using Guppy version 5.0.16 or 5.1.13. The Nanopore long reads were not cleaned and raw reads were used for genome assembly.

#### Illumina library preparation and sequencing

A PCR free library was prepared following the Kapa Hyper Prep Kit procedure provided by Roche (Roche, Basel, Switzerland). The library was sequenced on a paired-end mode with an Illumina NovaSeq 6000 instrument (Illumina, San Diego, CA, USA) using a 151 base-length read chemistry.

#### Hi-C library preparation and sequencing

Two Omni-C libraries were prepared using the Dovetail Omni-C Kit (Dovetail Genomics, Scotts Valley, CA, USA) following the Mammalian Cell Protocol for Sample Preparation version 1.4 after plant nuclei isolation. Briefly, flash-frozen young leaves (400 mg to 1 g) were cryoground in liquid nitrogen and pure nuclei were first isolated following the “High Molecular Weight DNA Extraction from Recalcitrant Plant Species’’ protocol described by Workman et al. (https://doi.org/10.1038/protex.2018.059). Once the nuclei had been isolated, the pellets were treated as mammalian cells and were fixed with DSG and formaldehyde; chromatin was randomly digested with DNase I and then extracted. Chromatin ends were repaired and ligated to a biotinylated bridge adapter, followed by proximity-ligation of adapter-containing ends. After proximity ligation, crosslinks were reversed and DNA was purified from proteins. Purified DNA was treated to remove biotin that was not internal to ligated fragments, and a sequencing library was generated using NEBNext Ultra enzymes and Illumina-compatible adapters. Biotin-containing fragments were isolated using streptavidin beads before PCR enrichment of the library. The final libraries were sequenced on an Illumina NovaSeq6000 instrument (Illumina, San Diego, CA, USA) with 2x 150 read length, and 83 million Omni-C reads were generated.

#### RNA library preparation and sequencing

Starting with 100 ng of total RNA, rRNA were first depleted from RNA samples extracted from leaves using the QIAseq FastSelect –rRNA Plant Kit (Qiagen, Hilde, Germany). A library was then prepared following the NEBNext Ultra II Directional RNA Library Prep for Illumina Kit procedure (New England Biolabs, Ipswich, MA, USA). A library was directly prepared from 100 ng of total RNA extracted from floral buds using the NEBNext Ultra II Directional RNA Library Prep for Illumina Kit. Libraries were sequenced with an Illumina NovaSeq 6000 instrument (Illumina) using a paired-end 151 base-length read chemistry.

#### Assembly of the *Arabidopsis halleri* reference genome

We generated three sets of read samples: the complete set of reads, 30X coverage of the longest reads, and 30X coverage of the filtlong (https://github.com/rrwick/Filtlong) highest-score reads (Supplemental Table S1). We then launched three different assemblers, Smartdenovo (Liu et al., 2021), Flye (Kolmogorov et al., 2019), and NECAT (Chen et al., 2021) on these three subsets of reads with the exception that NECAT was specifically run on the entire set of reads due to the implementation of a downsampling algorithm in its pipeline. Smartdenovo was launched with the parameters -k 17, as advised by the developers in case of larger genomes and -c 1 to generate a consensus sequence. Flye was launched with an estimated genome size of 240 Mbp and the -nano-raw option. NECAT was launched with a genome size of 240 Mbp and all other parameters set to their default values. Out of the 7 different assemblies obtained (Supplemental Table S8), we selected the Necat output for its higher contiguity (N50 > 1Mb) to continue our workflow. The Necat output was polished one time using Racon (Vaser et al., 2017) with Nanopore reads, then one time with Medaka (https://github.com/nanoporetech/medaka) (model r941_prom_hac_g507) and Nanopore reads, and two times with Hapo-G (1.3.4) (Aury and Istace, 2021) and Illumina short reads. We obtained an assembly of 518 contigs.

However the cumulative size of the assembly was higher than expected due to the high heterozygosity rate (320 Mb vs 240 Mb), and suggesting that the assembly size was currently inflated by the presence of allelic duplications. As indicated by BUSCO (Waterhouse et al., 2018) and KAT (Mapleson et al., 2017) (Supplemental Table S2 and Figure S9A), we observed the two alleles for many genes and a significant proportion of homozygous kmers were present twice in the assembly. We used HaploMerger2 (Huang et al., 2017) with default parameters and generated a haploid version of the assembly (Batch A twice to remove major misjoin and one run of Batch B). Haplomerger2 detected allelic duplications through all-against-all alignments and chose for each alignment the longest genomic regions (parameter - selectLongHaplotype), which may generate haplotype switches but ensure to maximize the gene content. We obtained two haplotypes: a reference version composed of the longer haplotype (when two haplotypes are available for a genomic locus) and a second version, named alternative, with the corresponding other allele of each duplicated genomic locus. At the end of the process, *A. halleri* haploid assembly has a cumulative size of 225 Mb, closer to the expected one, and KAT analysis showed a reduction of allelic duplications (Figure S9B). Additionally, the contig N50 benefited greatly from the separation and combination of the two haplotypes, rising to 3.3 Mb (Supplemental Table S2). Final assembly was polished one last time with Hapo-G and Illumina short reads to ensure that no allelic regions present twice in the diploid assembly have remained unpolished.

Chromosome-scale assembly was achieved using Hi-C data (Supplemental Table S1) and the 3D-DNA pipeline (version 180419) (https://github.com/aidenlab/3d-dna). Hi-C raw reads were aligned against the assembly (-s none option) using Juicer (Durand et al., 2016). The resulting merged_nodups.txt file and the assembly were given to the run-asm-pipeline.sh script with the options “--editor-repeat-coverage 5 -- splitter-coarse-stringency 30 --editor-coarse-resolution 100,000”. Contact maps were visualized through the Juicebox tool (version 1.11.08) (https://github.com/aidenlab/Juicebox) and edited to adjust the construction of chromosomes or break misjoins (Supplemental Figure S1). After edition, the new.assembly file was downloaded from the Juicebox interface, filtered and converted into a fasta file using the juicebox_assembly_converter.py script. Finally, Hapo-G was run one last time on the chromosome-scale haploid assembly.

#### *G*enome annotation of the *Arabidopsis halleri* reference assembly

The *A. halleri* reference genome was masked using RepeatMasker (v.4.1.0, default parameters) (Smit AFA, Hubley R, Green P. RepeatMasker. http://repeatmasker.org/) and a home-made library of transposable elements (based on four *Arabidopsis* species) available on the *A. halleri* repository (see Data availability section). Using this procedure, 48.6% of the input assembly was masked.

Gene prediction was done using as input homologous proteins and RNA-Seq data. Proteins from *Arabidopsis thaliana* (TAIR10) and *Arabidopsis lyrata* (extracted from uniprot database) were aligned against *A. halleri* masked genome assembly in two steps. Firstly, BLAT (default parameters) (Kent, 2002) was used to quickly localize corresponding putative genes of the proteins on the genome. The best match and matches with a score ≥ 90% of the best match score were retained. Secondly, the alignments were refined using Genewise (default parameters) (Birney et al., 2004), which is more precise for intron/exon boundary detection. Alignments were kept if more than 50% of the length of the protein was aligned to the genome.

To allow the detection of expressed and/or specific genes, we also used short-read RNA-Seq data extracted from two tissues (leaves and flower buds) of the same *A. halleri* individual. Short-reads were mapped on the genome assembly using HiSat2 (version 2.2.1 with default parameters) (Kim et al., 2019). Bam files were then sorted and merged by tissue and Stringtie (version 2.2.1) (Shumate et al., 2022) was launched on each tissue with the following parameters (--rf -p 16 -v -m 150). At each genomic locus, we kept only the most expressed transcript.

Finally, we integrated the protein homologies and transcripts using a combiner called Gmove (-m 10000 -e 3 -score) (Dubarry et al., 2016). This tool can find CDSs based on genome located evidence without any calibration step. Briefly, putative exons and introns, extracted from the alignments, were used to build a simplified graph by removing redundancies. Then, Gmove extracted all paths from the graph and searched for open reading frames (ORFs) consistent with the protein evidence. Completeness of the gene catalogs was assessed using BUSCO version 4.0.2 with the Brassica dataset odb10 and default parameters (Supplemental Table S2).

### Identification of miRNAs

#### sRNA extraction, library preparation and sequencing

Total RNA from *A. halleri,* Auby1, PL22, I30 and *A. lyrata* CP99 and MN47 samples were extracted with the miRNeasy minikit (Qiagen). For *A. halleri* PL22, I30 and *A. lyrata* CP99 and MN47, 3 µg of total RNA were sent to LC Sciences for library construction and sequencing. For *A. halleri* Auby1, total RNA (3µg) was purified with RNA Clean and Concentrator-5 kit (Zymo Research, Irvine, CA, USA, Ref. ZR1016), keeping only small RNAs fragments (17 - 200 nt) for the small RNAseq library preparation. Libraries were prepared following the NEXTflex Small RNA-Seq Kit v3 protocol provided by Perkin Elmer (Perkin Elmer, Waltham, MA, USA), starting with 200 ng of treated RNA, and ending with gel-free size selection and clean-up. The libraries were sequenced with an Illumina NovaSeq 6000 instrument (Illumina) using a paired-end 151 base-length read chemistry.

#### Additional data collection

Alongside the sRNA sequencing data produced in this study, various sets of sRNA sequencing data for *A. halleri*, *A. lyrata*, *A. thaliana, Camelina sativa, Capsella rubella, Raphanus sativus, Brassica oleracea, B. rapa, B. napus, B. juncea, B. nigra* and *Eutrema salsugineum* were obtained from the NCBI SRA database (https://www-ncbi-nlm-nih-gov.inee.bib.cnrs.fr/sra) (detailed information can be found in Supplemental Tables S4 and S7).

#### Identification of putative miRNA genes

The raw sRNA reads were processed according to miRkwood recommendations (https://bioinfo.univ-lille.fr/mirkwood/smallRNAseq/BED_file.php) using Python scripts performing adapter removal, trimming and quality filtering. Then, the sRNA reads were aligned to the reference genome of the respective species using Bowtie1 (Langmead et al., 2009), allowing for zero mismatch for the sample from *A. halleri* Auby1 and allowing for one mismatch for the other samples to be able detect isomirs (miRNA variants). The reference genomes used were those of the Auby1 (this present study) and MN47 accessions (Kolesnikova et al., 2023) for *A. halleri* and *A. lyrata*, respectively. For the remaining species, genome assemblies were downloaded from NCBI ASSEMBLY database (https://www-ncbi-nlm-nih-gov.inee.bib.cnrs.fr/assembly/) (detailed information can be found in Supplemental Table S5).

Our annotation strategy consisted of combining miRNAs predicted by miRkwood (score >=5) (Guigon et al., 2019) and Shortstack 4.0.2 (Johnson et al., 2016). miRkwood include a set of filters defined in Axtell and Meyers, (2018) such as a threshold for the stability of the hairpin (MFEI < -0.8), for the reads mapping to each arm of the hairpin (at least ten), the accuracy of precursor cleavage, the existence of the mature miRNA (read frequency at least 33%), the presence of the miRNA/miRNA* duplex and its stability (Guignon et al., 2019). Then, we merged the common predictions between the different samples and removed the predictions that fell into small chromosomal contigs to obtain a unique repertory for each species. Finally, to gain higher confidence in these predictions we mapped our sRNA read data onto predicted miRNA precursors using structVis v0.4 for manual observation (https://github.com/MikeAxtell/strucVis) (Supplemental Data Set S2).

### Experimental validation of miRNA predictions

#### Deep-sequencing of Argonaute-associated small RNAs

The Argonaute immunoprecipitations have been done according to Barre-Villeneuve et al. (2024) with anti-AGO1 antibodies (AS09 527, Agrisera) and Anti-AGO4 antibodies (AS09 617, Agrisera). Inflorescence, leaf and root tissues from pooled individuals of *A. halleri* (Auby and I9) and *A. lyrata* (Plech) were ground in liquid nitrogen and were homogenized in extraction buffer EB (50 mM Tris-HCl at pH 7.5, 150 mM NaCl, 5 mM MgCl_2_, 0.2% v/v NP40, 10% glycerol, 10 µM MG132) containing the EDTA-free protease inhibitor cocktail (Roche). After 15 min of incubation at 4°C, cell debris were removed by centrifugation at 21,000 *g* for 30 min at 4°C. The clarified lysate was incubated for 2 h at 4°C at 7-10 rpm, with AGO1 or AGO4 antibodies (from agrisera), and then 1 h at 4°C at 7-10 rpm with dynabeads protein A (Invitrogen), equilibrated with the EB. Beads were isolated using a magnetic rack, and washed once with 1 mL of EB and 4 times with 1 mL of PBS (Gibco). The sRNA were extracted from total/inputs and immunoprecipitated fractions using respectively Trizol and Trizol-LS, according to supplier instructions (Invitrogen). Subsequent sRNA libraries were performed and sequenced by the POPS platform from the plants science institute of Paris-Saclay (IPS2).

#### Bioinformatic analysis of AGO-IP libraries

After removal of adaptors, trimming and quality filtering, sequences were aligned onto the *A. halleri* and *A. lyrata* reference genomes with Bowtie1 allowing for one mismatch. We searched for an exact match between mature miRNA and sRNA read sequences and considered a miRNA loaded in AGO protein if more than 5 reads were found in the immunoprecipitate data. For each sample, reads were normalized per million total mapped reads (RPM). Enrichment with respect to the immunoprecipitate was calculated as the ratio of reads in the immunoprecipitate to reads in the input.

### Conservation analysis of miRNA genes

#### Synteny analysis of miRNA genes

The orthology maps of genes between *A. halleri* vs. *A. lyrata*, *A. halleri* vs. *A. thaliana* and *A. lyrata* vs. *A. thaliana* were constructed using protein sequences with OrthoFinder v2.5.4 (Emms and Kelly, 2019) using default parameters. Only orthogroups that contain one-to-one orthologues per species were kept for further comparison. Orthologous miRNAs between *A. halleri*, *A. lyrata* and *A. thaliana* were identified using the gene orthology maps described above. We selected miRNA genes located between upstream and downstream orthologous genes and restricted the size of the chromosomal fragment to 100 kb. The sequences of framed miRNA genes were aligned using the best-hit approach, commonly used to establish orthology relationships within genomes (Ward and Moreno-Hagelsieb, 2014). Two miRNA genes were considered syntenic if they were a reciprocal best match.

#### miRNA genes conservation across Viridiplantae

The miRNA families and the conservation across Viridiplantae were assigned based on similarity of mature miRNA sequences using the PmiREN 1.0 database (Guo et al., 2020). This database is specialized for plant miRNAs and is based on a standardized analysis of sRNA-seq data, which reduces the variability between predictions that would be due to the use of different tools. We filtered the database on mature miRNA sequence length requiring 18-nt to 25-nt sequences. In addition, we enriched the database with the predicted miRNAs from ten Brassicaceae species (*A. thaliana, Brassica juncea, B. napus, B. nigra, B. oleracea, B. rapa, Capsella rubella, Eutrema salsugineum, Camelina sativa, Raphanus sativus*), allowing us to be more precise about the conservation status of the miRNAs inside the Brassicaceae family. Then, the sequences of the mature miRNAs were aligned using Exonerate (Slater et al., 2005), allowing for three mismatch/gap/insertion. Alignments with a unique distant species (outside the Brassicaceae family) were considered as false positives.

#### Characterization of features of miRNA genes

We assessed the thermodynamic stability of the precursors using the Minimum Free Energy Index (MFEI) according to the equation MFEI = [MFE / sequence length x 100] / (G+C%) (Guignon et al., 2019). We determined the secondary structure MFE of the precursors using the RNAfold software (Lorenz et al., 2011) and used Python scripts to calculate the GC content.

From secondary structure, we further defined the different parts of the miRNA precursors (miRNA/miRNA* duplex, loop, stem and the flanking regions) using python scripts.

We calculated the abundance miRNAs in each sample where they were predicted and took the average value. The abundance of precursors was defined as the reads mapping the precursor normalized per 1,000,000 total mapped reads and the precursor length (RPKM). The abundance of mature miRNAs was normalized per 1,000,000 total mapped reads (RPM).

The associations between miRNA features and their age were examined with regression linear models using R (v4.1.2; R Core Team 2023).

### Target characterization

#### Target prediction

We identified the potential miRNA targets in the CDS of *A. halleri* and *A. lyrata* using TargetFinder (Bo and Wang, 2005) with default parameters, which provide the best balance between specificity and sensitivity (Srivastava et al., 2014). We applied a cut-off penalty score of ≤ 3 as recommended in Fahlgren et al., (2007) for reliable miRNA-mRNA target interactions.

#### Proxies of essentiality of *A. halleri* and *A. lyrata* genes

Three proxies have been used as in Legrand et al., (2019) to assess gene essentiality. Briefly, the all-against-all Blast method was employed using the CDS to estimate the size of the gene family. The hits with a query coverage inferior to 50% and/or an e-value superior to 1e-30 were discarded. Ka/Ks was estimated using KaKs_Calculator2.0 (Wang et al., 2010) with the Goldman and Yang method (Goldman et al., 1994) from the alignments of pairs of orthologous CDS between *A. halleri* vs. *A. thaliana* and *A. lyrata* vs. *A. thaliana* obtained using Water from the EMBOSS package (Rice et al., 2000). Finally, loss of function genes were identified using a dataset composed of 2400 Arabidopsis genes with a loss-of-function mutant phenotype (Lloyd and Meinke, 2012).

The associations between target gene features and miRNA gene ages were examined with regression linear models using R (v4.1.2; R Core Team 2023), except for loss-of-function genes proxy for which we used a Chi-squared test on all conservation groups.

### Polymorphism analysis

#### Data collection

To assess the genomic diversity, we analyzed 100 *A. lyrata* individuals from natural accessions. In addition to the genomic data produced, we downloaded WGS data obtained by Takou et al., (2021) and Mattila et al., (2017). The set was composed of 39 individuals from Michigan, USA (this study); Spiterstulen, Norway (24 individuals) (Mattila et al., 2017; Takou et al., 2021); Stubbsand, Sweden (6 individuals) (Mattila et al., 2017); Plech, Germany (18 individuals) (Mattila et al., 2017; Takou et al., 2021); Austria (7 individuals) (Takou et al., 2021); Mayodan, USA (6 individuals) (Mattila et al., 2017).

#### Variant calling and pi calculation

After adapters removal, the reads were mapped to the reference genomes of *A. halleri* and *A. lyrata* using bowtie2 (Langmead et al., 2012) and PCR duplicated reads were removed with picard MarkDuplicates version 2.21.4 (available at http://broadinstitute.github.io/picard). GATK version 4.1.9.0 (McKenna et al., 2010) was used to call and annotate single nucleotide polymorphisms (SNPs) using haplotypecaller. Individual GVCF files were subjected to joint genotyping to obtain a .vcf file with information on all sites, both variant and invariant. We extracted the precursors, targets and flanking regions and filtered the resulting .vcf files with VCFtools version 0.1.16 (Danecek et al., 2011). Because invariant sites do not have quality scores, we created individual .vcf files for variant and invariant sites. Invariant sites were identified by setting the minor allele frequency to zero (--max-maf), whereas variant sites have a minor allele count ≥1 (--mac). We filtered variant sites using the following options --remove-indels --min-alleles 1 --max-alleles 2 --max-missing 0.75 --minDP 5 --minQ 30. Subsequently, we indexed both .vcf files with tabix of SAMtools and combined them with BCFtools version 1.12 (Danecek et al., 2021). The average number of nucleotide differences between genotypes (π) was calculated using VCFtools version 0.1.16 (Danecek et al., 2011). The average number of nucleotide differences between genotypes (π) was calculated using VCFtools version 0.1.16 (Danecek et al., 2011). Additionally, we carried out permutation tests to assess the probability that the differences we observed could be due to our result being different from chance alone and thus determining its significance. Specifically, For example, each nucleotide associated with its nucleotide diversity value (π) was permuted within the hairpin, and then the average π of each region of the hairpin was calculated. This was repeated a large number of times (*n*=1000), allowing us to define a confidence interval. If the average π observed for the hairpin part was outside the confidence interval, this meant that the observed value was different from chance and therefore significant.

## Acknowledgments

We thank the high performance computing service and Bilille at the University of Lille for providing computing resources. This work was performed using the infrastructure and technical support of the “Plateforme Serre, cultures et terrains expérimentaux – Université de Lille” for the greenhouse/field facilities. We thank Anamaria Nesculesca, Filipe Borges for taking part in FP’s PhD committee, and Blake Meyers, Noah Fahlgren, Patricia Baldrich and Xavier Vekemans for discussions.

## Availability of supporting data

The Illumina and Oxford Nanopore sequencing data of the *A. halleri* reference genome are available in the European Nucleotide Archive under the following project PRJEB70878. The small RNA sequencing data are available in the NCBI-SRA database under the following BioProject PRJNA1098478. All the previously undiscovered miRNA loci were deposited at miRBase (https://www.mirbase.org; Kozomara et al. 2019). The supplementary data sets are available in Figshare under the following accession numbers : https://doi.org/10.6084/m9.figshare.25746267.v1, https://doi.org/10.6084/m9.figshare.25737294.v3 and https://doi.org/10.6084/m9.figshare.25737288.v3

## Credit authorship contribution statement

Cultivated plants: CPo, EH, FP. Performed molecular biology experiments: FP, JAF, CB, CC, LD, RAB, VK. Analysed data : FP, SL, CPa, FL, SG, JMA, EL, RAB, MG. Contributed samples: VK, UK. Designed the project : VC, SL, JAF, ED. Wrote the manuscript : FP, VC, SL, JMA, EL.

## References

Allen, E., Xie, Z., Gustafson, A.M., Sung, G.-H., Spatafora, J.W., and Carrington, J.C. (2004). Evolution of microRNA genes by inverted duplication of target gene sequences in Arabidopsis thaliana. Nat Genet 36: 1282–1290.

Aury, J.-M. and Istace, B. (2021). Hapo-G, haplotype-aware polishing of genome assemblies with accurate reads. NAR Genomics and Bioinformatics 3: lqab034.

Axtell, M.J. and Meyers, B.C. (2018). Revisiting Criteria for Plant MicroRNA Annotation in the Era of Big Data. Plant Cell 30: 272–284.

Baldrich, P., Beric, A., and Meyers, B.C. (2018). Despacito: the slow evolutionary changes in plant microRNAs. Current Opinion in Plant Biology 42: 16–22.

Barre-Villeneuve, C., Laudié, M., Carpentier, M.-C., Kuhn, L., Lagrange, T., and Azevedo-Favory, J. (2024). The unique dual targeting of AGO1 by two types of PRMT enzymes promotes phasiRNA loading in *Arabidopsis thaliana*. Nucleic Acids Research: gkae045.

Belser, C. et al. (2018). Chromosome-scale assemblies of plant genomes using nanopore long reads and optical maps. Nature Plants 4: 879–887.

Bénitìere, F., Necsulea, A., and Duret, L. (2024). Random genetic drift sets an upper limit on mRNA splicing accuracy in metazoans.

Birney, E., Clamp, M., and Durbin, R. (2004). GeneWise and Genomewise. Genome Res. 14: 988–995.

Bo, X. and Wang, S. (2005). TargetFinder: a software for antisense oligonucleotide target site selection based on MAST and secondary structures of target mRNA. Bioinformatics 21: 1401–1402.

Bologna, N.G., Schapire, A.L., Zhai, J., Chorostecki, U., Boisbouvier, J., Meyers, B.C., and Palatnik, J.F. (2013). Multiple RNA recognition patterns during microRNA biogenesis in plants. Genome Res. 23: 1675–1689.

Borges, F., Parent, J.-S., Van Ex, F., Wolff, P., Martínez, G., Köhler, C., and Martienssen, R.A. (2018). Transposon-derived small RNAs triggered by miR845 mediate genome dosage response in Arabidopsis. Nat Genet 50: 186–192.

Bradley, D. et al. (2017). Evolution of flower color pattern through selection on regulatory small RNAs. Science 358: 925–928.

Briskine, R.V., Paape, T., Shimizu-Inatsugi, R., Nishiyama, T., Akama, S., Sese, J., and Shimizu, K.K. (2017). Genome assembly and annotation of *Arabidopsis halleri*, a model for heavy metal hyperaccumulation and evolutionary ecology. Molecular Ecology Resources 17: 1025–1036.

Carvunis, A.-R. et al. (2012). Proto-genes and de novo gene birth. Nature 487: 370– 374.

Chávez Montes, R.A., Rosas-Cárdenas, D.F.F., De Paoli, E., Accerbi, M., Rymarquis, L.A., Mahalingam, G., Marsch-Martínez, N., Meyers, B.C., Green, P.J., and De Folter, S. (2014). Sample sequencing of vascular plants demonstrates widespread conservation and divergence of microRNAs. Nat Commun 5: 3722.

Chen, K. and Rajewsky, N. (2007). The evolution of gene regulation by transcription factors and microRNAs. Nat Rev Genet 8: 93–103.

Chen, Y. et al. (2021). Efficient assembly of nanopore reads via highly accurate and intact error correction. Nat Commun 12: 60.

Crescente, J.M., Zavallo, D., Del Vas, M., Asurmendi, S., Helguera, M., Fernandez, E., and Vanzetti, L.S. (2022). Genome-wide identification of MITE-derived microRNAs and their targets in bread wheat. BMC Genomics 23: 154.

Cui, J., You, C., and Chen, X. (2017). The evolution of microRNAs in plants. Current Opinion in Plant Biology 35: 61–67.

Cuperus, J.T., Fahlgren, N., and Carrington, J.C. (2011). Evolution and Functional Diversification of *MIRNA* Genes. Plant Cell 23: 431–442.

Danecek, P., Auton A, Abecasis G, Albers CA, Banks E, DePristo MA, Handsaker RE, Lunter G, Marth GT, Sherry ST, McVean G, Durbin R; 1000 Genomes Project Analysis Group (2011). The variant call format and VCFtools. Bioinformatics 27: 2156–2158.

Danecek P., Bonfield J.K., Liddle J., Marshall J., Ohan V., Pollard M.O., Whitwham A., Keane T., McCarthy S.A., Davies R.M., and Li H. (2021) Twelve years of SAMtools and BCFtools. GigaScience, 10, 2021, 1–4

Dexheimer, P.J. and Cochella, L. (2020). MicroRNAs: From Mechanism to Organism. Front. Cell Dev. Biol. 8: 409.

Ding, N. and Zhang, B. (2023). microRNA production in Arabidopsis. Front. Plant Sci. 14: 1096772.

Dong, Q., Hu, B., and Zhang, C. (2022). microRNAs and Their Roles in Plant Development. Front. Plant Sci. 13: 824240.

Dubarry, M. et al., Gmove a Tool for Eukaryotic Gene Predictions Using Various Evidences (F1000Research, 2016).

Durand, E., Méheust, R., Soucaze, M., Goubet, P.M., Gallina, S., Poux, C., Fobis-Loisy, I., Guillon, E., Gaude, T., Sarazin, A., Figeac, M., Prat, E., Marande, W., Bergès, H., Vekemans, X., Billiard, S., Castric, V. (2014). Dominance hierarchy arising from the evolution of a complex small RNA regulatory network. Science 346: 1200–1205.

Durand, N.C., Shamim, M.S., Machol, I., Rao, S.S.P., Huntley, M.H., Lander, E.S., and Aiden, E.L. (2016). Juicer Provides a One-Click System for Analyzing Loop-Resolution Hi-C Experiments. Cell Systems 3: 95–98.

Erdmann R.M. and Picard C.L. RNA-directed DNA Methylation. PLoS Genet. 2020:16(10):e1009034. 10.1371/journal.pgen.1009034

Ehrenreich, I.M. and Purugganan, M.D. (2008). Sequence Variation of MicroRNAs and Their Binding Sites in Arabidopsis. Plant Physiology 146: 1974–1982.

Emms, D.M. and Kelly, S. (2019). OrthoFinder: phylogenetic orthology inference for comparative genomics. Genome Biol 20: 238.

Fahlgren, N., Howell, M.D., Kasschau, K.D., Chapman, E.J., Sullivan, C.M., Cumbie, J.S., Givan, S.A., Law, T.F., Grant, S.R., Dangl, J.L., and Carrington, J.C. (2007). High-Throughput Sequencing of Arabidopsis microRNAs: Evidence for Frequent Birth and Death of MIRNA Genes. PLoS ONE 2: e219.

Fahlgren, N., Sullivan, C.M., Kasschau, K.D., Chapman, E.J., Cumbie, J.S., Montgomery, T.A., Gilbert, S.D., Dasenko, M., Backman, T.W.H., Givan, S.A., and Carrington, J.C. (2009). Computational and analytical framework for small RNA profiling by high-throughput sequencing. RNA 15: 992–1002.

Fahlgren, N., Jogdeo, S., Kasschau, K.D., Sullivan, C.M., Chapman, E.J., Laubinger, S., Smith, L.M., Dasenko, M., Givan, S.A., Weigel, D., and Carrington, J.C. (2010). MicroRNA Gene Evolution in *Arabidopsis lyrata* and *Arabidopsis thaliana*. The Plant Cell 22: 1074–1089.

François Jacob (1977). Evolution and Tinkering. Science 196: 1661–1666.

Goldman N. and Yang Z. (1994) A codon-based model of nucleotide substitution for protein-coding DNA sequences. Molecular Biology and Evolution.

Goudarzi, M., Berg, K., Pieper, L.M., and Schier, A.F. (2019). Individual long non-coding RNAs have no overt functions in zebrafish embryogenesis, viability and fertility. eLife 8: e40815.

Guigon, I., Legrand, S., Berthelot, J.-F., Bini, S., Lanselle, D., Benmounah, M., and Touzet, H. (2019). miRkwood: a tool for the reliable identification of microRNAs in plant genomes. BMC Genomics 20: 532.

Guo, Z. et al. (2020). PmiREN: a comprehensive encyclopedia of plant miRNAs. Nucleic Acids Research 48: D1114–D1121.

Guo, Z., Kuang, Z., Deng, Y., Li, L., and Yang, X. (2022a). Identification of Species-Specific MicroRNAs Provides Insights into Dynamic Evolution of MicroRNAs in Plants. IJMS 23: 14273.

Guo, Z., Kuang, Z., Zhao, Y., Deng, Y., He, H., Wan, M., Tao, Y., Wang, D., Wei, J., Li, L., and Yang, X. (2022b). PmiREN2.0: from data annotation to functional exploration of plant microRNAs. Nucleic Acids Research 50: D1475–D1482.

Huang, S., Kang, M., and Xu, A. (2017). HaploMerger2: rebuilding both haploid sub-assemblies from high-heterozygosity diploid genome assembly. Bioinformatics 33: 2577–2579.

Kent, W.J. (2002) BLAT—The BLAST-Like Alignment Tool.

Kim, D., Paggi, J.M., Park, C., Bennett, C., and Salzberg, S.L. (2019). Graph-based genome alignment and genotyping with HISAT2 and HISAT-genotype. Nat Biotechnol 37: 907–915.

Koch, M.A., Haubold, B., and Mitchell-Olds, T. (2000). Comparative Evolutionary Analysis of Chalcone Synthase and Alcohol Dehydrogenase Loci in Arabidopsis, Arabis, and Related Genera (Brassicaceae). Molecular Biology and Evolution 17: 1483–1498.

Kolesnikova, U.K., Scott, A.D., Van De Velde, J.D., Burns, R., Tikhomirov, N.P., Pfordt, U., Clarke, A.C., Yant, L., Seregin, A.P., Vekemans, X., Laurent, S., and Novikova, P.Y. (2023). Transition to Self-compatibility Associated With Dominant *S* -allele in a Diploid Siberian Progenitor of Allotetraploid *Arabidopsis kamchatica* Revealed by *Arabidopsis lyrata* Genomes. Molecular Biology and Evolution 40: msad122.

Kolmogorov, M., Yuan, J., Lin, Y., and Pevzner, P.A. (2019). Assembly of long, error-prone reads using repeat graphs. Nat Biotechnol 37: 540–546.

Kozomara, A., Birgaoanu, M., and Griffiths-Jones, S. (2019). miRBase: from microRNA sequences to function. Nucleic Acids Research 47: D155–D162.

Kubota, S., Iwasaki, T., Hanada, K., Nagano, A.J., Fujiyama, A., Toyoda, A., Sugano, S., Suzuki, Y., Hikosaka, K., Ito, M., and Morinaga, S.-I. (2015). A Genome Scan for Genes Underlying Microgeographic-Scale Local Adaptation in a Wild Arabidopsis Species. PLoS Genet 11: e1005361.

Kumar, S., Suleski, M., Craig, J.M., Kasprowicz, A.E., Sanderford, M., Li, M., Stecher, G., and Hedges, S.B. (2022). TimeTree 5: An Expanded Resource for Species Divergence Times. Molecular Biology and Evolution 39: msac174.

Johnson, N.R., Yeoh, J.M., Coruh, C., and Axtell, M.J. Improved Placement of Multi-mapping Small RNAs. G3 Genes|Genomes|Genetics 6:2103–2111.

Langmead, B., Trapnell, C., Pop, M., and Salzberg, S.L. (2009). Ultrafast and memory-efficient alignment of short DNA sequences to the human genome. Genome Biol 10: R25.

Langmead, B. and Salzberg, S.L. (2012). Fast gapped-read alignment with Bowtie 2. Nat Methods 9: 357–359.

Legrand, S. et al. (2019). Differential retention of transposable element-derived sequences in outcrossing Arabidopsis genomes. Mobile DNA 10: 30.

Li, J., Reichel, M., Li, Y., and Millar, A.A. (2014). The functional scope of plant microRNA-mediated silencing. Trends in Plant Science 19: 750–756.

Li, Q., Liu, G., Bao, Y., Wu, Y., and You, Q. (2021). Evaluation and application of tools for the identification of known microRNAs in plants. Appl Plant Sci 9.

Liu, H., Wu, S., Li, A., and Ruan, J. (2021). SMARTdenovo: a de novo assembler using long noisy reads. Gigabyte 2021: 1–9.

Lloyd, J. and Meinke, D. (2012). A Comprehensive Dataset of Genes with a Loss-of-Function Mutant Phenotype in Arabidopsis. Plant Physiology 158: 1115–1129.

Lorenz, R., Bernhart, S.H., Höner Zu Siederdissen, C., Tafer, H., Flamm, C., Stadler, P.F., and Hofacker, I.L. (2011). ViennaRNA Package 2.0. Algorithms Mol Biol 6: 26.

Lynch, M. (2007). The evolution of genetic networks by non-adaptive processes. Nat Rev Genet 8: 803–813.

Ma, Z., Coruh, C., and Axtell, M.J. (2010). *Arabidopsis lyrata* Small RNAs: Transient *MIRNA* and Small Interfering RNA Loci within the *Arabidopsis* Genus. Plant Cell 22: 1090–1103.

Mapleson, D., Garcia Accinelli, G., Kettleborough, G., Wright, J., and Clavijo, B.J. (2017). KAT: a K-mer analysis toolkit to quality control NGS datasets and genome assemblies. Bioinformatics 33: 574–576.

Mattila, T.M., Tyrmi, J., Pyhäjärvi, T., and Savolainen, O. (2017). Genome-Wide Analysis of Colonization History and Concomitant Selection in Arabidopsis lyrata. Molecular Biology and Evolution 34: 2665–2677.

McLysaght, A. and Guerzoni, D. (2015). New genes from non-coding sequence: the role of de novo protein-coding genes in eukaryotic evolutionary innovation. Phil. Trans. R. Soc. B 370: 20140332.

Mi, S. et al. (2008). Sorting of Small RNAs into Arabidopsis Argonaute Complexes Is Directed by the 5′ Terminal Nucleotide. Cell 133: 116–127.

Nanbo, A., Furuyama, W., and Lin, Z. (2021). RNA Virus-Encoded miRNAs: Current Insights and Future Challenges. Front. Microbiol. 12: 679210.

Nozawa, M., Fujimi, M., Iwamoto, C., Onizuka, K., Fukuda, N., Ikeo, K., and Gojobori, T. (2016). Evolutionary Transitions of MicroRNA-Target Pairs. Genome Biol Evol 8: 1621–1633.

Nozawa, M., Miura, S., and Nei, M. (2012). Origins and Evolution of MicroRNA Genes in Plant Species. Genome Biology and Evolution 4: 230–239.

Ossowski, S., Schneeberger, K., Lucas-Lledó, J.I., Warthmann, N., Clark, R.M., Shaw, R.G., Weigel, D., and Lynch, M. (2010). The Rate and Molecular Spectrum of Spontaneous Mutations in *Arabidopsis thaliana*. Science 327: 92– 94.

Pegler, J.L., Oultram, J.M.J., Mann, C.W.G., Carroll, B.J., Grof, C.P.L., and Eamens, A.L. (2023). Miniature Inverted-Repeat Transposable Elements: Small DNA Transposons That Have Contributed to Plant MICRORNA Gene Evolution. Plants 12: 1101.

Poretti, M., Praz, C.R., Meile, L., Kälin, C., Schaefer, L.K., Schläfli, M., Widrig, V., Sanchez-Vallet, A., Wicker, T., and Bourras, S. (2020). Domestication of High-Copy Transposons Underlays the Wheat Small RNA Response to an Obligate Pathogen. Molecular Biology and Evolution 37: 839–848.

Rajagopalan, R., Vaucheret, H., Trejo, J., and Bartel, D.P. (2006). A diverse and evolutionarily fluid set of microRNAs in *Arabidopsis thaliana*. Genes Dev. 20: 3407–3425.

Rice, P. EMBOSS: The European Molecular Biology Open Software Suite.

Roux, C., Castric, V., Pauwels, M., Wright, S.I., Saumitou-Laprade, P., and Vekemans, X. (2011). Does Speciation between Arabidopsis halleri and Arabidopsis lyrata Coincide with Major Changes in a Molecular Target of Adaptation? PLoS ONE 6: e26872.

Roux, J., Gonzàlez-Porta, M., and Robinson-Rechavi, M. (2012). Comparative analysis of human and mouse expression data illuminates tissue-specific evolutionary patterns of miRNAs. Nucleic Acids Research 40: 5890–5900.

Shumate, A., Wong, B., Pertea, G., and Pertea, M. (2022). Improved transcriptome assembly using a hybrid of long and short reads with StringTie. PLoS Comput Biol 18: e1009730.

Slater, G. and Birney, E. (2005). Automated generation of heuristics for biological sequence comparison. BMC Bioinformatics 6: 31.

Statello, L., Guo, C.-J., Chen, L.-L., and Huarte, M. (2021). Gene regulation by long non-coding RNAs and its biological functions. Nat Rev Mol Cell Biol 22: 96– 118.

Smith, L.M., Burbano, H.A., Wang, X., Fitz, J., Wang, G., Ural-Blimke, Y., and Weigel, D. (2015). Rapid divergence and high diversity of miRNAs and miRNA targets in the Camelineae. Plant J 81: 597–610.

Srivastava, P.K., Moturu, T.R., Pandey, P., Baldwin, I.T., and Pandey, S.P. (2014). A comparison of performance of plant miRNA target prediction tools and the characterization of features for genome-wide target prediction. BMC Genomics 15: 348.

Takou, M., Hämälä, T., Koch, E.M., Steige, K.A., Dittberner, H., Yant, L., Genete, M., Sunyaev, S., Castric, V., Vekemans, X., Savolainen, O., and Meaux, J.D. (2021). Maintenance of Adaptive Dynamics and No Detectable Load in a Range-Edge Outcrossing Plant Population. Molecular Biology and Evolution 38: 1820– 1836.

Vacherie B., Labadie, K., Falentin, C., (2022). HMW DNA extraction for Long Read Sequencing using CTAB. protocols.io 10.17504/protoc ols.io.bp2l694yzlqe/v1

Van Oss, S.B. and Carvunis, A.-R. (2019). De novo gene birth. PLoS Genet 15: e1008160.

Vaser, R., Sović, I., Nagarajan, N., and Šikić, M. (2017). Fast and accurate de novo genome assembly from long uncorrected reads. Genome Res. 27: 737– 746.

Voinnet, O. (2009). Origin, Biogenesis, and Activity of Plant MicroRNAs. Cell 136: 669–687.

Wagner, A. (2011). The molecular origins of evolutionary innovations. Trends in Genetics 27: 397–410.

Wang, D., Zhang, Y., Zhang, Z., Zhu, J., and Yu, J. (2010). KaKs_Calculator 2.0: A Toolkit Incorporating Gamma-Series Methods and Sliding Window Strategies. Genomics, Proteomics & Bioinformatics 8: 77–80.

Ward, N. and Moreno-Hagelsieb, G. (2014). Quickly Finding Orthologs as Reciprocal Best Hits with BLAT, LAST, and UBLAST: How Much Do We Miss? PLoS ONE 9: e101850.

Waterhouse, R.M., Seppey, M., Simão, F.A., Manni, M., Ioannidis, P., Klioutchnikov, G., Kriventseva, E.V., and Zdobnov, E.M. (2018). BUSCO Applications from Quality Assessments to Gene Prediction and Phylogenomics. Molecular Biology and Evolution 35: 543–548.

Wong, G.Y. and Millar, A.A. (2023). Target Landscape of Conserved Plant MicroRNAs and the Complexities of Their Ancient MicroRNA-Binding Sites. Plant Cell Physiol 64:604–621.

Wen, M., Lin, X., Xie, M., Wang, Y., Shen, X., Liufu, Z., Wu, C.-I., Shi, S., and Tang, T. (2016). Small RNA transcriptomes of mangroves evolve adaptively in extreme environments. Sci Rep 6: 27551.

Wilson, B.A., Foy, S.G., Neme, R., and Masel, J. (2017). Young genes are highly disordered as predicted by the preadaptation hypothesis of de novo gene birth. Nat Ecol Evol 1: 0146.

Wright, C., Rajpurohit, A., Burke, E.E., Williams, C., Collado-Torres, L., Kimos, M., Brandon, N.J., Cross, A.J., Jaffe, A.E., Weinberger, D.R., and Shin, J.H. (2019). Comprehensive assessment of multiple biases in small RNA sequencing reveals significant differences in the performance of widely used methods. BMC Genomics 20: 513.

Zhan, J. and Meyers, B.C. (2023). Plant Small RNAs: Their Biogenesis, Regulatory Roles, and Functions. Annu. Rev. Plant Biol. 74: 21–51.

Zhang Y, Jiang W, and Gao L. Evolution of MicroRNA Genes in Oryza sativa and Arabidopsis thaliana: An Update of the Inverted Duplication Model. PLoS ONE. 2011:6(12):e28073.

