## Supplemental data for "The evolutionary history and functional specialization of microRNA genes in *Arabidopsis halleri* and *A. lyrata*"

**Supplementary data**

**Table S1. Comparison of nanopore readset statistics.**

| | Full | Filtlong | Longest |
|---|---|---|---|
| Number of Bases (GB) | 29 | 7.2 | 7.2 |
| Coverage | 122 | 30 | 30 |
| Number of Reads | 3 322 114 | 230 775 | 173 520 |
| N50 | 18 929 | 33 132 | 40 619 |
| Average Size | 8 854 | 31 199 | 41 494 |

**Table S2. Assembly statistics of the reference accession (Auby1) throughout the process.**

| | Necat | Polished | Haplomerger2 | Chromosome Scale |
|---|---|---|---|---|
| Total length (Mb) | 323 | 323 | 227 | 227 |
| Number of scaffolds/contigs | 509 | 518 | 284 | 175 |
| N50 (Kb) | 1 597 | 1 600 | 3 347 | 25 922 |
| L50 | 57 | 57 | 20 | 5 |
| Average contig size (Kb) | 634 | 625 | 800 | 1 298 |
| Merqury score | 24.4648 | 31.8868 | 32.1278 | 32.7199 |
| Complete universal single-copy orthologs | C:98.4% S:59.4% D:39.0% | C:99.3% S:57.9% D:41.4% | C:99.0% S:97.0% D:2.0% | C:99.1% S:97.2% D:1.9% |
| Fragmented universal single-copy orthologs | 0.6% | 0.2 | 0.3% | 0.2% |
| Missing universal single-copy orthologs | 1.0% | 0.5% | 0.7% | 0.7% |

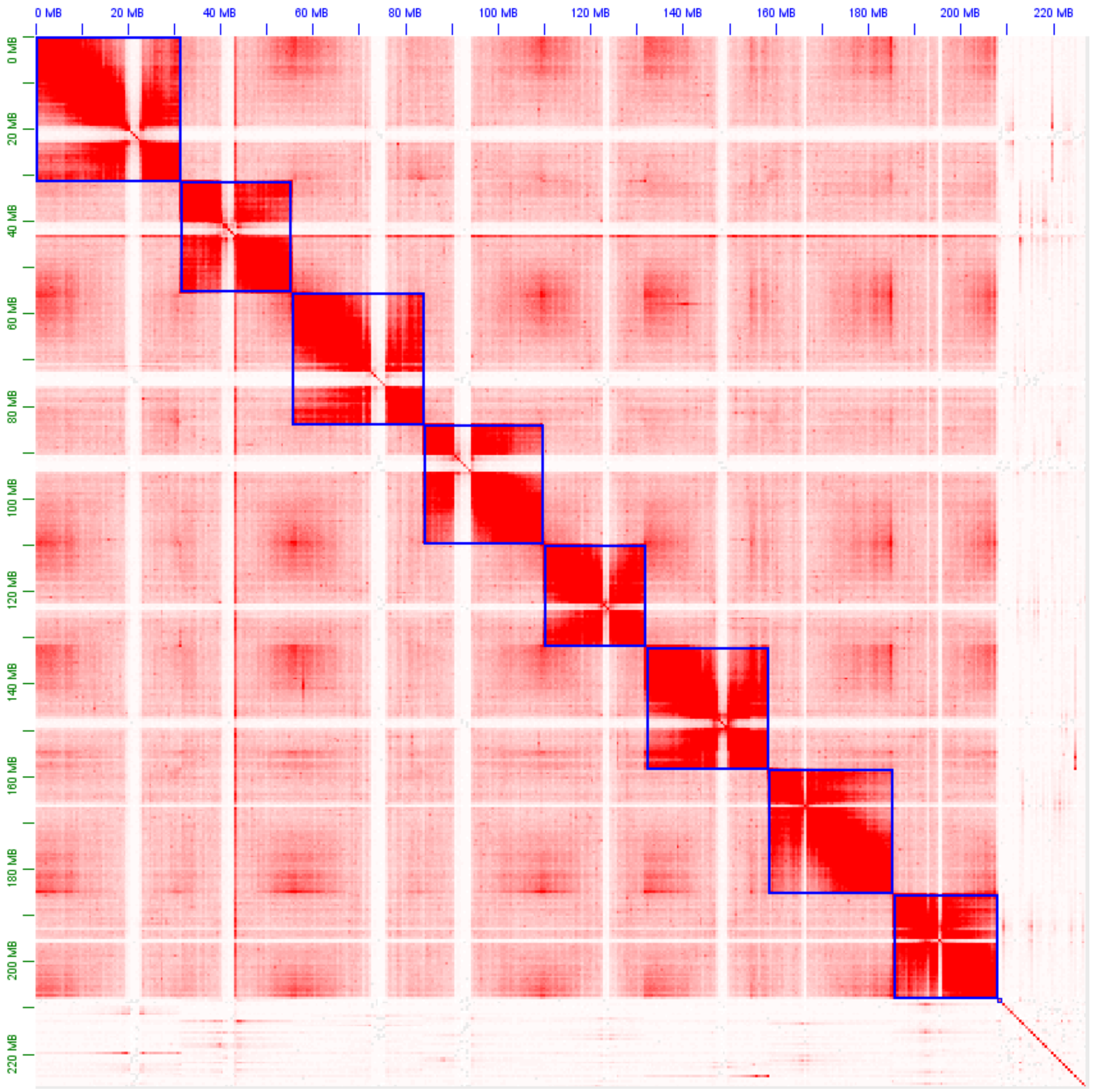


**Figure S1. Curated chromosome-scale assembly of a reference *A. halleri* accession (Auby-1).** The red dots correspond to Hi-C contacts. The green squares correspond to contigs from the PACBIO assembly, and are assembled into chromosome-level scaffolds represented by blue squares.

**Table S3. Comparison of *A. halleri* PL22 and Auby1 genome assemblies.**

| Genome assembly metrics | *A. halleri* PL22 (Legrand et al. 2019) | *A. halleri* Auby1 (this study) |
|---|---|---|
| Number of contigs/ scaffolds | 3152 | 175 |
| Total length (Mb) | 174 | 227 |
| N50 | 279,389 | 25,922,902 |
| L50 | 177 | 5 |
| Longest contig/scaffold (Mb) | 1.5 | 31 |
| Complete universal single-copy orthologs | 95.3% | 99.1% |
| Fragmented universal single-copy orthologs | 1.5% | 0.2% |
| Missing universal single-copy orthologs | 3.2% | 0.7% |

**Table S4. sRNAseq datasets for miRNA predictions.**

| Species | Accession | Tissue | Library preparation | Sequencing technology | Total reads[a] | Number of miRNA genes[b] | Reference | SRA-NCBI |
|---|---|---|---|---|---|---|---|---|
| *A.halleri* | Auby1 | Leaves | Nextflex® Small RNA-Seq | Illumina | 206,983,903 | 196 | This study | PRJEB70878 |
| *A.halleri* | Auby1 | Buds | Nextflex® Small RNA-Seq | Illumina | 159,726,202 | 267 | This study | PRJEB70878 |
| *A.halleri* | Auby | Roots | Nextflex® Small RNA-Seq | Illumina | 41,470,720 | 80 | This study | SRR28676106 |
| *A.halleri* | Auby | Buds | Nextflex® Small RNA-Seq | Illumina | 40,304,810 | 121 | This study | SRR28676098 |
| *A.halleri* | Auby | Leaves | Nextflex® Small RNA-Seq | Illumina | 40,236,011 | 72 | This study | SRR28676096 |
| *A.halleri* | I9 | Roots | Nextflex® Small RNA-Seq | Illumina | 40,564,542 | 71 | This study | SRR28676105 |
| *A.halleri* | I9 | Buds | Nextflex® Small RNA-Seq | Illumina | 39,370,067 | 100 | This study | SRR28676109 |
| *A.halleri* | I9 | Leaves | Nextflex® Small RNA-Seq | Illumina | 41,102,882 | 84 | This study | SRR28676102 |
| *A.halleri* | PL22 | Leave | Nextflex® Small RNA-Seq | Illumina | 9,403,700 | 124 | This study | SRR28676112 |
| *A.halleri* | I30 | Leaves | Nextflex® Small RNA-Seq | Illumina | 13,304,075 | 109 | This study | SRR28676111 |
| *A.halleri* | HF70 | Buds | SOLiD Total RNA-Seq | SOLiD | 31,926,829 | 74 | Durand et al. (2014) | SRR1271746 |
| *A.halleri* | I5 | Buds | SOLiD Total RNA-Seq | SOLiD | 27,201,005 | 91 | Durand et al. (2014) | SRR1271747 |
| *A.halleri* | I5 | Buds | SOLiD Total RNA-Seq | SOLiD | 28,166,676 | 96 | Durand et al. (2014) | SRR1271748 |
| *A.halleri* | I5 | Buds | SOLiD Total RNA-Seq | SOLiD | 27,521,772 | 79 | Durand et al. (2014) | SRR1271749 |
| *A.halleri* | I9 | Buds | SOLiD Total RNA-Seq | SOLiD | 27,760,235 | 77 | Durand et al. (2014) | SRR1271750 |
| *A.halleri* | Nivelle | Buds | SOLiD Total RNA-Seq | SOLiD | 24,590,979 | 82 | Durand et al. (2014) | SRR1271751 |
| *A.halleri* | Nivelle | Buds | SOLiD Total RNA-Seq | SOLiD | 26,216,099 | 69 | Durand et al. (2014) | SRR1271752 |
| *A.halleri* | Nivelle | Buds | SOLiD Total RNA-Seq | SOLiD | 30,136,006 | 77 | Durand et al. (2014) | SRR1271753 |
| *A.halleri* | BC01 | Buds | ION total RNA-seq | Proton | 23,441,621 | 99 | Durand et al. (2014) | SRR1271755 |
| *A.halleri* | BC02 | Buds | ION total RNA-seq | Proton | 21,907,916 | 98 | Durand et al. (2014) | SRR1271756 |
| *A.halleri* | BC03 | Buds | ION total RNA-seq | Proton | 23,779,085 | 93 | Durand et al. (2014) | SRR1271757 |
| *A.lyrata* | MN47 | Leaves | TruSeq Small RNA | Illumina | 9,012,391 | 102 | Legrand et al. (2019) | SRR9665460 |

| | | | | | | | | |
|---|---|---|---|---|---|---|---|---|
| *A.lyrata* | CP99 | Leaves | NA | Illumina | 28,995,456 | 95 | This study | SRR28676084 |
| *A.lyrata* | CP99 | Buds | NA | Illumina | 10,424,454 | 73 | This study | SRR28676091 |
| *A.lyrata* | Al14 | Buds | SOLiD Total RNA-Seq | SOLiD | 36,006,064 | 69 | Durand et al. (2014) | SRR1271754 |
| *A.lyrata* | MN47 | Leaves | Ma et al. (2010) | Illumina | 2,012,409 | 46 | Ma et al. (2010) | SRR034856 |
| *A.lyrata* | MN47 | Buds | SOLiD Small RNA Expression | SOLiD | 90,518,311 | 126 | Ma et al. (2010) | SRR040401 |
| *A.lyrata* | MN47 | Buds | SOLiD Small RNA Expression | SOLiD | 10,321,920 | 67 | Ma et al. (2010) | SRR040402 |
| *A.lyrata* | MN47 | Leaves | Fahlgren et al. (2009) | Illumina | 5,093,642 | 62 | Fahlgren et al. (2010) | SRR051926 |
| *A.lyrata* | MN47 | Buds | Fahlgren et al. (2010) | Illumina | 4,876,824 | 61 | Fahlgren et al. (2010) | GSM518430[c] |
| *A.lyrata* | MN47 | Buds | Fahlgren et al. (2010) | Illumina | 4,229,395 | 64 | Fahlgren et al. (2010) | GSM518431[c] |
| *A.lyrata* | Plech | Roots | Nextflex® Small RNA-Seq | Illumina | 43,024,185 | 64 | This study | SRR28676090 |
| *A.lyrata* | Plech | Buds | Nextflex® Small RNA-Seq | Illumina | 41,899,496 | 128 | This study | SRR28676094 |
| *A.lyrata* | Plech | Leaves | Nextflex® Small RNA-Seq | Illumina | 36,480,074 | 110 | This study | SRR28676087 |

[a] Total number of reads in the sequencing experiment.
[b] Number of miRNA genes predicted in the sequencing experiment.
[c] Data in gff3 format while all the others are in fastq format.

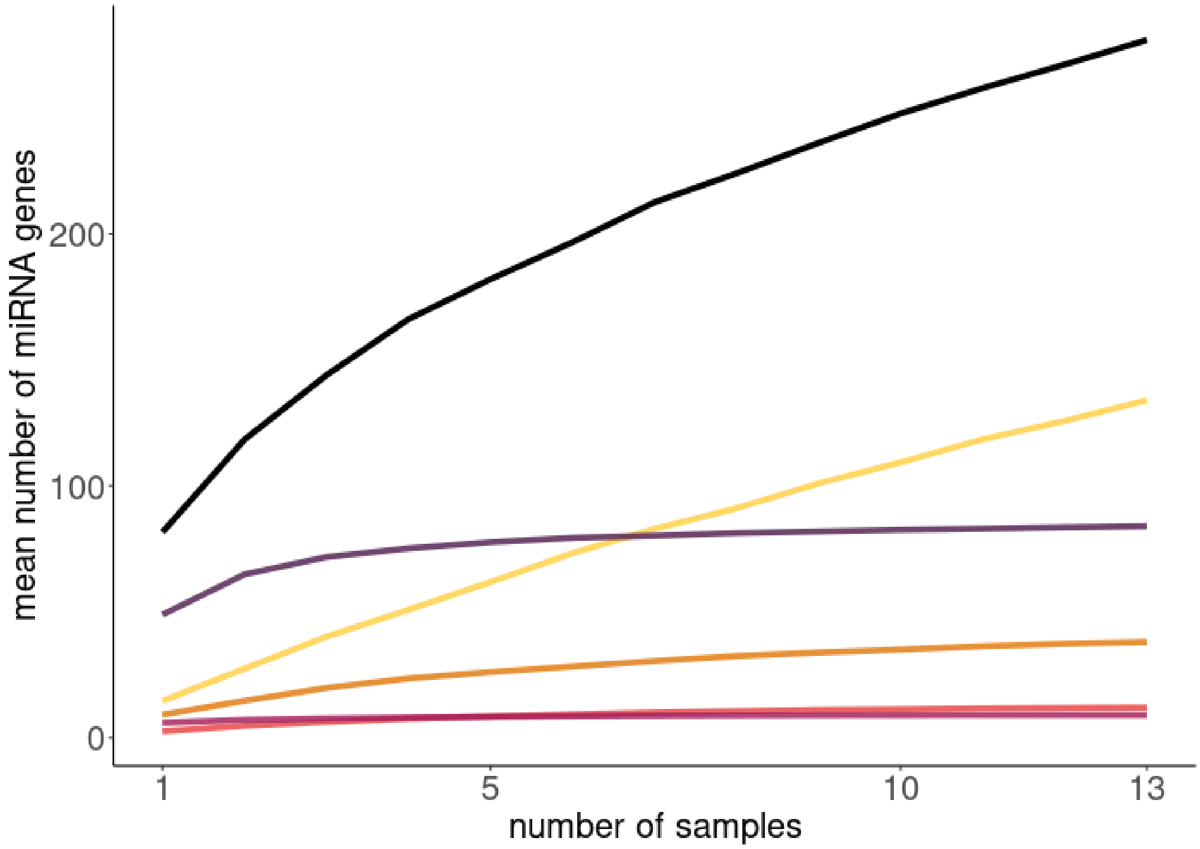


**Figure S2. Completeness of the miRNA gene repertories according to the numbers of individuals sampled in *A. lyrata*.** The black line indicates the entire repertoire, the purple line the repertoire of conserved miRNA genes, the pink line, the repertoire of miRNA genes shared by Brassicaceae species, the dark orange line, the ones shared by *A. thaliana*, *A. halleri* and *A. lyrata*, the light orange line, the miRNA genes shared between *A. halleri* and *A. lyrata* and the yellow line the ones specific to *A. lyrata*.

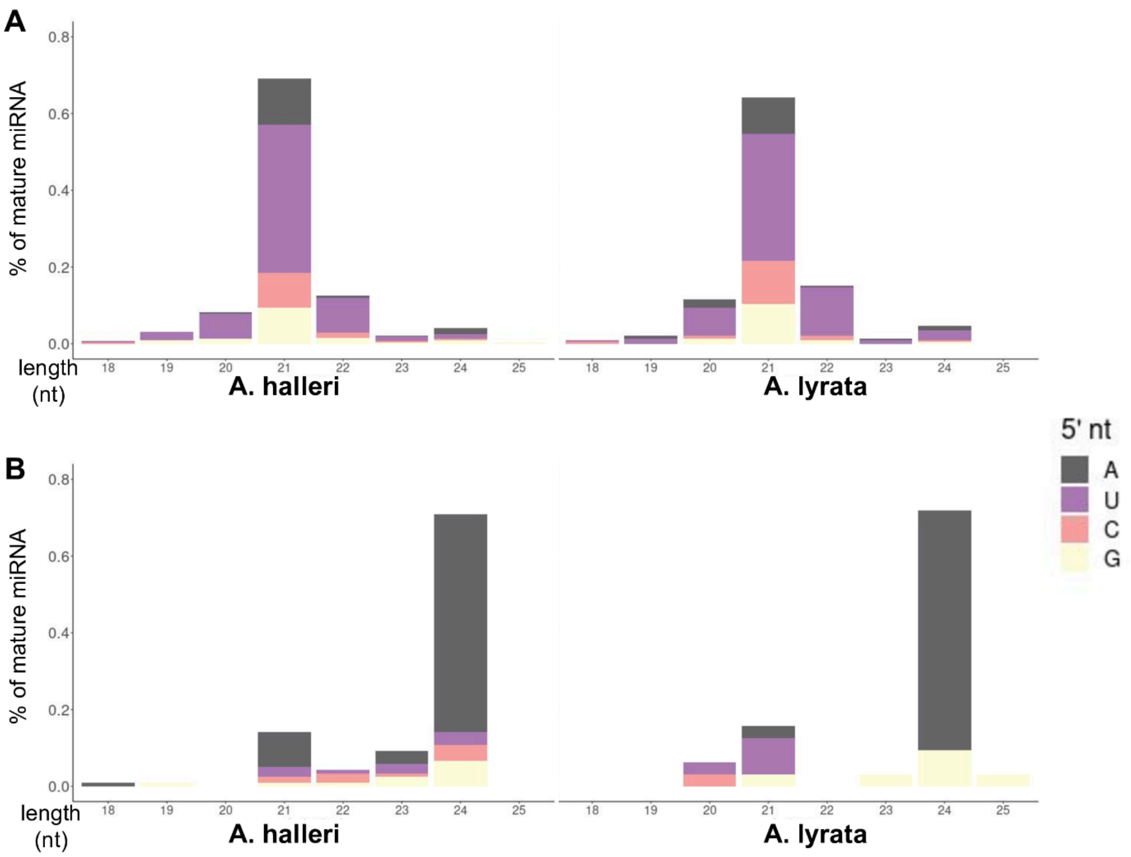


**Figure S3. Size distribution and nature of the 5’nt of AGO1 (a) and AGO4 (b) -associated miRNAs.**

**Table S5. sRNAseq Datasets for miRNA predictions in the Brassicaceae family.**

| Species | Tissu | Genome assembly version | Study for sRNAseq libraries | SRA-NCBI | Sequencing technology | Total reads number | Number of miRNA genes | Number of mature miRNA |
|---|---|---|---|---|---|---|---|---|
| *A. thaliana* | Leaves | TAIR10 | Wang et al. 2022 | SRR17231451 | Illumina | 9,747,585 | 127 | 145 |
| *A. thaliana* | Roots | TAIR10 | Blein et al. 2020 | SRR8723396 | Proton | 15,832,411 | 160 | 199 |
| *A. thaliana* | Seedlings | TAIR10 | Choi et al. 2021 | SRR15082674 | Illumina | 87,270,697 | 122 | 154 |
| *A. thaliana* | Seedlings | TAIR10 | Choi et al. 2021 | SRR15082675 | Illumina | 98,587,809 | 138 | 182 |
| *A. thaliana* | Seedlings | TAIR10 | Choi et al. 2021 | SRR15082676 | Illumina | 67,931,288 | 106 | 135 |
| *A. thaliana* | Buds | TAIR10 | in prep | SRR27110143 | Illumina | 31,307,342 | 130 | 155 |
| *A. thaliana* | Buds | TAIR10 | in prep | SRR27110146 | Illumina | 55,852,923 | 162 | 190 |
| *A. thaliana* | Buds | TAIR10 | in prep | SRR28386575 | Illumina | 21,377,860 | 129 | 159 |
| *Camelina sativa* | Leaves | GCF_000633955.1_Cs | Poudel et al. 2015 | SRR1736515 | Illumina | 9,856,027 | 164 | 173 |
| *Capsella rubella* | Leaves | GCF_000375325.1_Caprub1_0 | Smith et al. 2014 | SRR942635 | Illumina | 24,194,069 | 118 | 126 |
| *Raphanus sativus* | Leaves | GCF_000801105.1_Rs1.0 | Yang et al. 2019 | SRR7725716 | Illumina | 15,239,556 | 152 | 182 |
| *Brassica oleracea* | Leaves | Boleraceacapitata_446_v1.0 | Lukasik et al. 2013 | SRR799357 | Illumina | 24,037,208 | 91 | 96 |
| *Brassica rapa* | Leaves | GCF_000309985.2_CAAS_Brap_v3.02 | Ahmed et al. 2020 | SRR11092574 | Illumina | 25,331,960 | 147 | 166 |
| *Brassica napus* | Leaves | GCF_000686985.2_Bra_napus_v2.0 | Regmi et al. 2021 | SRR13071038 | Illumina | 22,377,118 | 164 | 180 |
| *Brassica juncea* | Leaves | GCA_018703725.1_ASM1870372v1 | Cao et al. 2016 | SRR3441529 | Illumina | 12,365,840 | 156 | 184 |
| *Brassica nigra* | Leaves | GCA_016432835.1_Bnig_sang_1.1 | Ghani et al. 2014 | SRR1592476 | Illumina | 10,284,599 | 133 | 142 |
| *Eutrema salsugineum* | Leaves | GCF_000478725.1_Eutsalg1_0 | Niederhuth et al. 2016 | SRR3286330 | Illumina | 9,602,054 | 78 | 85 |

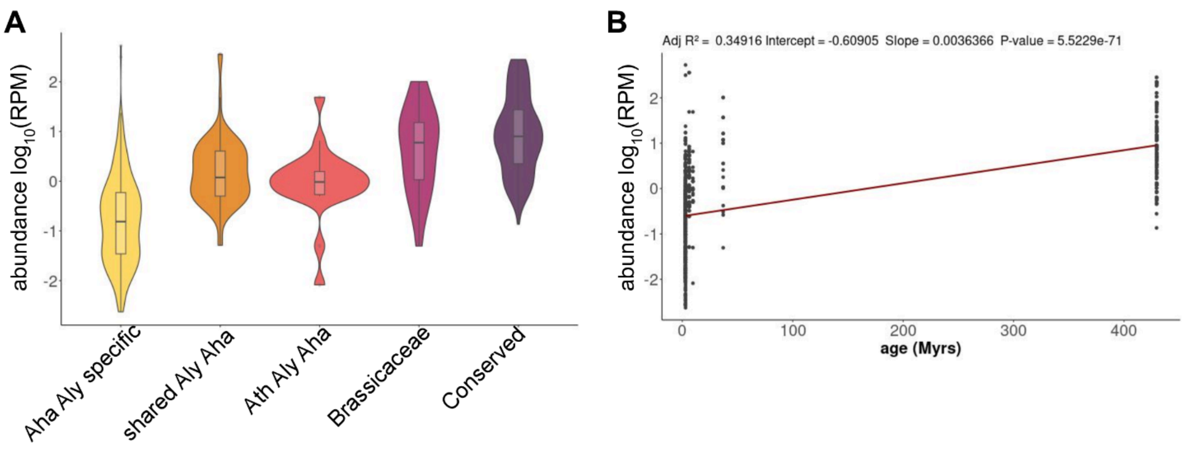


**Figure S4. Mature miRNA expression according to their conservation.** (a) mature miRNA abundance (RPM) according to age of the miRNA gene. (b) Linear regression of the mature miRNA expression according to the age.

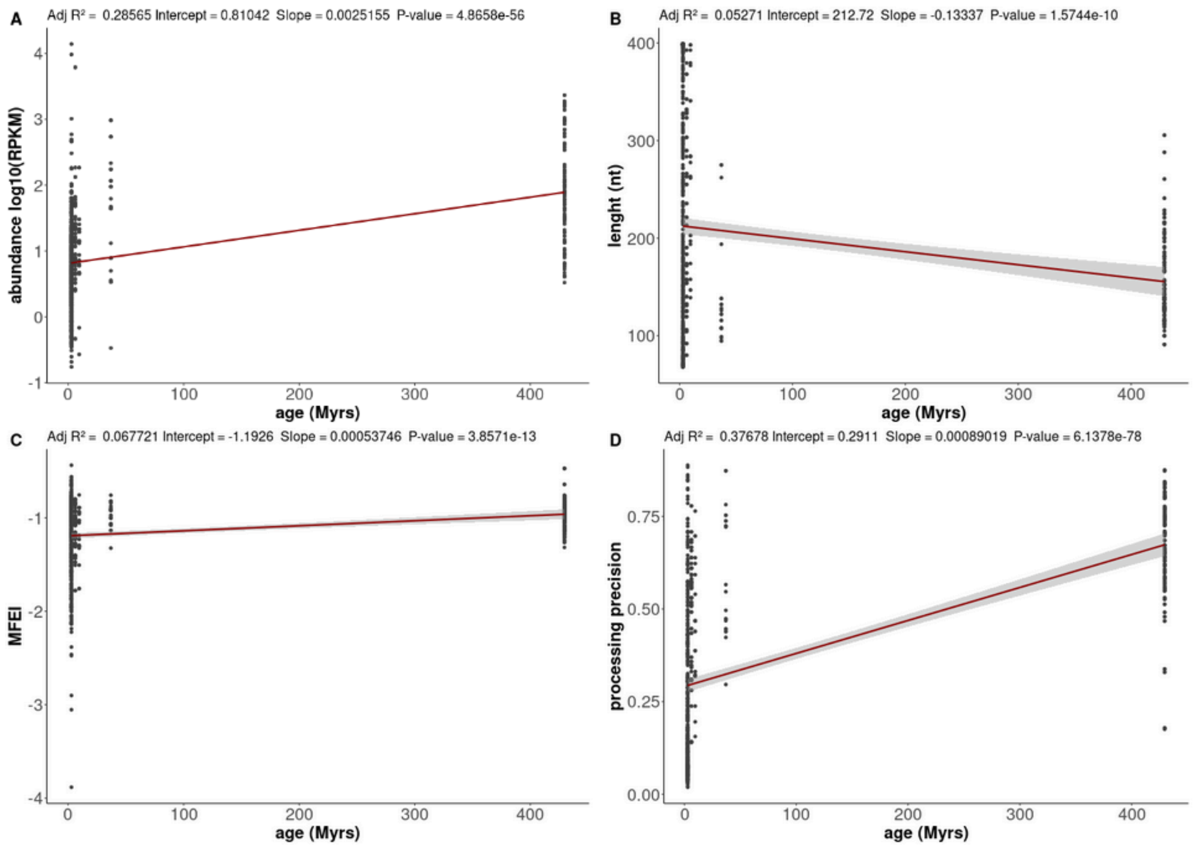


**Figure S5: Linear regression of the miRNA genes characteristics according to their age.** (a) precursor abundance (RPKM) according to age. (b) predicted hairpin length (c) hairpin stability (MFEI) (d) processing precision.

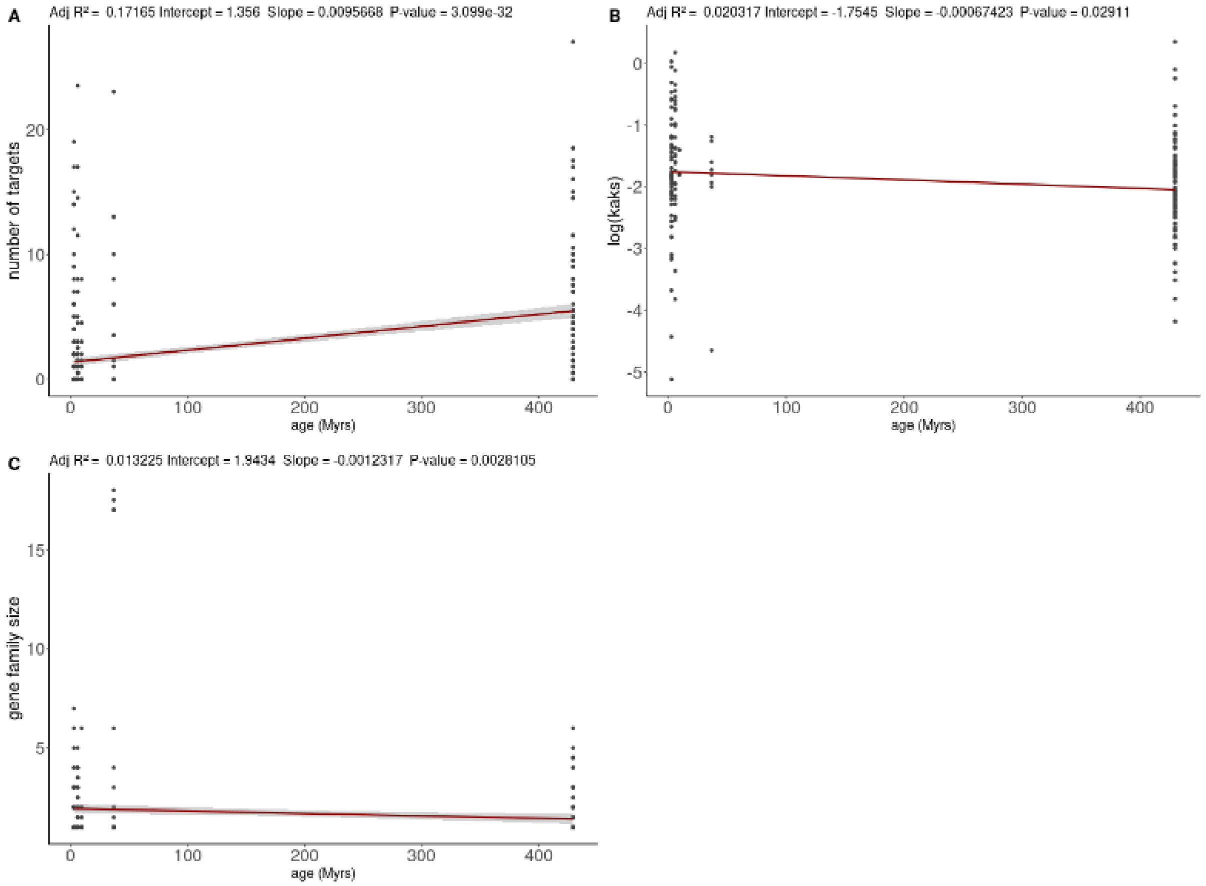


**Figure S6. Linear regression of the miRNA genes target characteristics according to their age.** (a) Number of targets (b) $k_A/k_S$ ratios (c) family size.

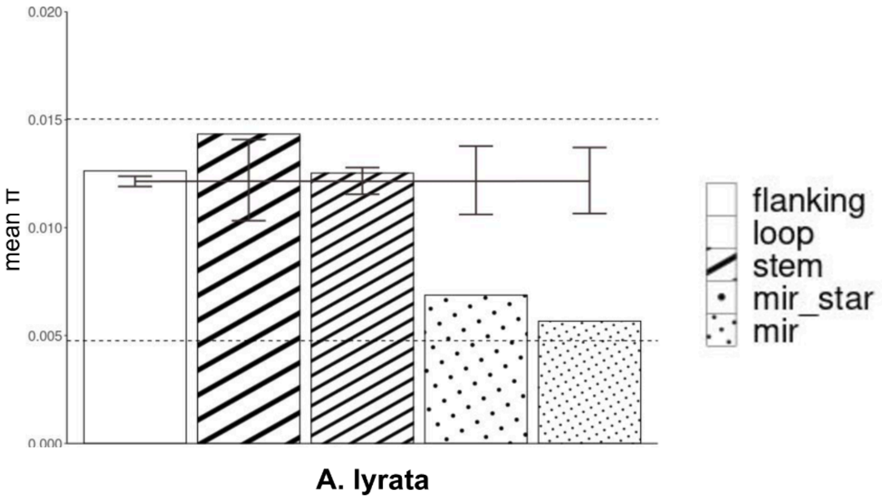


**Figure S7. The miRNA/miRNA* duplex is strongly constrained by natural selection.** Average nucleotide diversity for the different parts of the miRNA hairpins and upstream and downstream flanking regions (200 bp each) in *A. lyrata.* The dashed lines represent the mean π value for the 0 fold (lower bar) and 4 fold (upper bar) degenerate positions of all protein-coding genes. The bars represent the 95% confidence interval obtained by random permutation of nucleotides for 1,000 random permutations under the hypothesis of a uniform distribution of polymorphisms along the sequence.

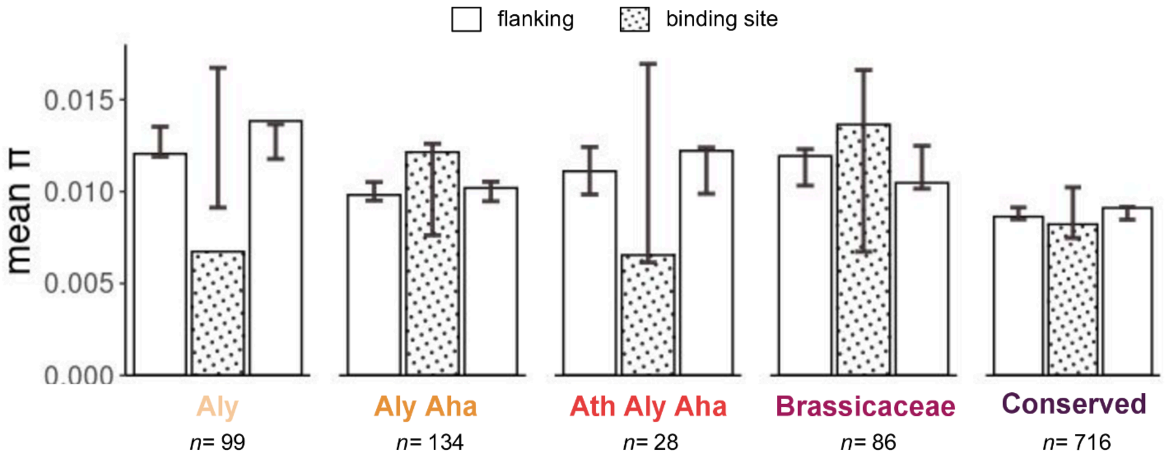


**Figure S8. Average nucleotide diversity for the miRNA binding site and upstream and downstream flanking regions (300 bp each) in *A. lyrata* mRNA targets according to the conservation of the miRNA gene targeting them*.***

The bars represent the 95% confidence interval obtained by random permutation of nucleotides for 1,000 simulations.

**Table S8.** Phylogenetic families of the 87 plant species used to analyze the conservation of the miRNA genes.

<table>
<tr><td colspan="5">Bryophytes</td><td>Physcomitrella patens</td></tr>
<tr><td rowspan="33">Tracheophytes</td><td rowspan="33">Spermatophytes</td><td colspan="3" rowspan="2">Spermatophytes</td><td>Picea abies</td></tr>
<tr><td>Ginkgo biloba</td></tr>
<tr><td rowspan="31">Magnoliophyta</td><td rowspan="18">Liliopsida</td><td>Araceae</td><td>Spirodela polyrhiza</td></tr>
<tr><td>Zosteraceae</td><td>Zostera marina</td></tr>
<tr><td>Orchidaceae</td><td>Phalaenopsis aphrodite</td></tr>
<tr><td>Asparagaceae</td><td>Asparagus officinalis</td></tr>
<tr><td>Arecaceae</td><td>Phoenix dactylifera</td></tr>
<tr><td rowspan="12">Poaceae</td><td>Zea mays</td></tr>
<tr><td>Saccharum hybrid</td></tr>
<tr><td>Sorghum bicolor</td></tr>
<tr><td>Panicum hallii</td></tr>
<tr><td>Panicum virgatum</td></tr>
<tr><td>Setaria italica</td></tr>
<tr><td>Brachypodium distachyon</td></tr>
<tr><td>Triticum aestivum</td></tr>
<tr><td>Oryza sativa</td></tr>
<tr><td>Oryza rufipogon</td></tr>
<tr><td>Oryza nivara</td></tr>
<tr><td>Oryza glaberrima</td></tr>
<tr><td>Musaceae</td><td>Musa acuminata</td></tr>
<tr><td rowspan="13">Eudicotyledones</td><td>Nelumbonaceae</td><td>Nelumbo nucifera</td></tr>
<tr><td rowspan="7">Fabaceae</td><td>Lotus japonicus</td></tr>
<tr><td>Medicago truncatula</td></tr>
<tr><td>Pisum sativum</td></tr>
<tr><td>Cicer arietinum</td></tr>
<tr><td>Glycine max</td></tr>
<tr><td>Phaseolus vulgaris</td></tr>
<tr><td>Vigna unguiculata</td></tr>
<tr><td rowspan="3">Rosaceae</td><td>Prunus persica</td></tr>
<tr><td>Fragaria ananassa</td></tr>
<tr><td>Fragaria vesca</td></tr>
<tr><td rowspan="2">Fagaceae</td><td>Fagus sylvatica</td></tr>
<tr><td>Quercus robur</td></tr>
</table>

| | | | | | |
|---|---|---|---|---|---|
| | | | | Cucurbitaceae | *Citrullus lanatus* |
| | | | | | *Lagenaria siceraria* |
| | | | | | *Cucumis melo* |
| | | | | | *Cucurbita maxima* |
| | | | | | *Cucurbita moschata* |
| | | | | Euphorbiaceae | *Ricinus communis* |
| | | | | | *Jatropha curcas* |
| | | | | | *Manihot esculenta* |
| | | | | | *Hevea brasiliensis* |
| | | | | Salicaceae | *Populus euphratica* |
| | | | | | *Populus trichocarpa* |
| | | | | Lythraceae | *Punica granatum* |
| | | | | Rutaceae | *Citrus clementina* |
| | | | | | *Citrus maxima* |
| | | | | | *Citrus sinensis* |
| | | | | Caricaceae | *Carica papaya* |
| | | | | Brassicaceae | *Arabidopsis thaliana* |
| | | | | | *Arabidopsis lyrata* |
| | | | | | *Arabidopsis halleri* |
| | | | | | *Camelina sativa* |
| | | | | | *Capsella rubella* |
| | | | | | *Raphinus sativus* |
| | | | | | *Brassica oleracea* |
| | | | | | *Brassica rapa* |
| | | | | | *Brassica napus* |
| | | | | | *Brassica juncea* |
| | | | | | *Brassica nigra* |
| | | | | | *Eutrema salsugineum* |
| | | | | Malvaceae | *Hibiscus syriacus* |
| | | | | | *Gossypium hirsutum* |
| | | | | | *Gossypium barbadense* |
| | | | | | *Gossypium arboreum* |
| | | | | Vitaceae | *Vitis vinifera* |
| | | | | Actinidiaceae | *Actinidia chinensis* |

| | | | | | |
|---|---|---|---|---|---|
| | | | | Asteraceae | *Carthamus tinctorius* |
| | | | | | *Lactuca sativa* |
| | | | | Solanaceae | *Nicotiana benthamiana* |
| | | | | | *Nicotiana tabacum* |
| | | | | | *Solanum melongena* |
| | | | | | *Solanum tuberosum* |
| | | | | | *Solanum pennellii* |
| | | | | | *Solanum pimpinellifolium* |
| | | | | | *Solanum lycopersicum* |
| | | | | | *Solanum habrochaites* |
| | | | | | *Capsicum annuum* |
| | | | | Convolvulaceae | *Ipomoea batatas* |
| | | | | Oleaceae | *Fraxinus excelsior* |
| | | | | | *Olea europaea* |
| | | | | Phrymaceae | *Erythranthe guttata* |
| | | | | Caryophyllaceae | *Silene latifolia* |
| | | | Magnoliidae | | *Persea americana* |
| | | | Magnoliophyta | | *Amborella trichopoda* |
| | Lycophytes | | | | *Selaginella moellendorffii* |

**Table S7. Sample GPS coordinates**

| Species | Accession name | GPS Coordinates |
|---|---|---|
| *A. halleri* | Auby (France) | 50.40400191 3.091988208 |
| *A. halleri* | PL22 (Poland) | 50.282800, 19.478717 |
| *A. halleri* | I9 (Italy) | 46.73141, 11.43292 |
| *A. halleri* | I30 (Italy) | 45.991119, 10.272050 |
| *A. lyrata* | Plech (Germany) | 49.627550, 11.511536 |
| *A. lyrata* | CP99 (Czech Republic) | 50.0854713, 14.1303275 |
| *A. lyrata* | LPT (USA) | -80.3875, 42.5797222 |
| *A. lyrata* | TC (USA) | -81.51750000000001, 45.2416667 |
| *A. lyrata* | TSS (USA) | -81.58388889999999, 45.1925 |
| *A. lyrata* | PIN (USA) | -81.8313889, 43.26888890000001 |
| *A. lyrata* | RON (USA) | -81.8463889, 42.2613889 |
| *A. lyrata* | IND (USA) | -87.0422175, 41.6689766 |

**Table S8. Comparison of assemblies statistics.**

| | Necat | SmartDenovo | | | Flye | | |
|---|---|---|---|---|---|---|---|
| Readset | Full | Full | Filtlong | Longest | Full | Filtlong | Longest |
| Total length (Mb) | 323 | 327 | 224 | 224 | 280 | 303 | 295 |
| Number of contigs | 509 | 1209 | 707 | 707 | 4 205 | 2 956 | 2 740 |
| N50 (Kb) | 1 597 | 871 | 770 | 770 | 204 | 231 | 254 |
| L50 | 57 | 86 | 64 | 64 | 347 | 332 | 292 |
| Average contig size (Kb) | 634 | 270 | 317 | 317 | 67 | 103 | 107 |
| Merqury score | 24.4648 | 23.0902 | 21.4396 | 19.8713 | 24.727 | 23.2474 | 23.208 |
| Complete universal single-copy orthologs | C:98.4% S:59.4% D:39.0% | C:98.5% S:76.9% D:21.6% | C:95.1% S:87.3% D:7.8% | C:93.0% S:87.0% D:6.0% | C:99.1% S:77.7% D:21.4% | C:99.0% S:76.8% D:22.2% | C:98.6% S:73.8% D:24.8% |
| Fragmented universal single-copy orthologs | 0.6% | 0.6% | 1.3% | 1.6%, | 0.5% | 0.5% | 0.6% |
| Missing universal single-copy orthologs | 1.0% | 0.9% | 3.6% | 5.4% | 0.4% | 0.5% | 0.5% |

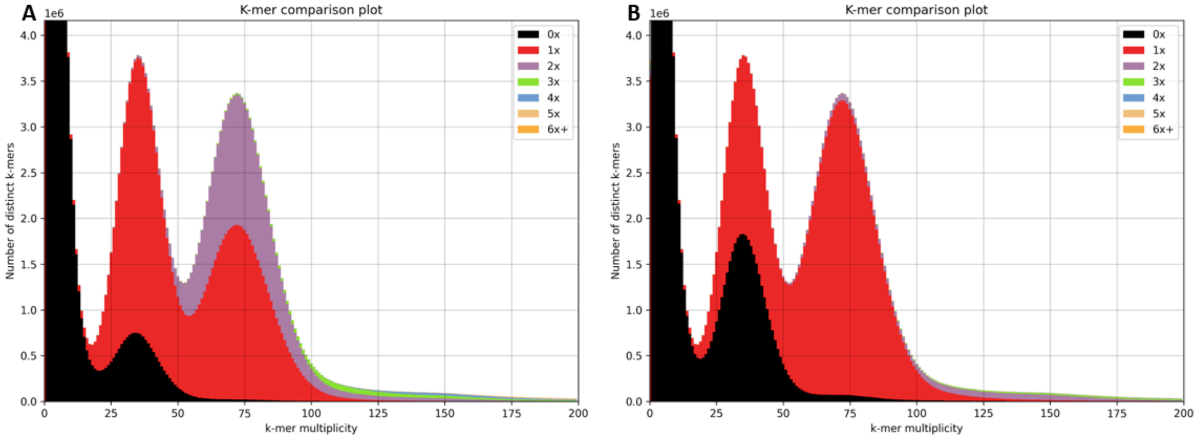


**Figure S9. KAT plot. (A) Pre-Haplomerger2. (B) Post-Haplomerger2**
